# Signature patterns for top-down and bottom-up information processing via cross-frequency coupling in macaque auditory cortex

**DOI:** 10.1101/403980

**Authors:** Christian D. Márton, Makoto Fukushima, Corrie R. Camalier, Simon R. Schultz, Bruno B. Averbeck

## Abstract

Predictive coding is a theoretical framework that provides a functional interpretation of top-down and bottom up interactions in sensory processing. The theory has suggested that specific frequency bands relay bottom-up and top-down information (e.g. “*γ* up, *β* down”). But it remains unclear whether this notion generalizes to cross-frequency interactions. Furthermore, most of the evidence so far comes from visual pathways. Here we examined cross-frequency coupling across four sectors of the auditory hierarchy in the macaque. We computed two measures of cross-frequency coupling, phase-amplitude coupling (PAC) and amplitude-amplitude coupling (AAC). Our findings revealed distinct patterns for bottom-up and top-down information processing among *cross*-frequency interactions. Both top-down and bottom-up made prominent use of low frequencies: low-to-low frequency (*θ, α, β*) and low frequency-to-*high γ* couplings were predominant *top-down*, while low frequency-to-*low γ* couplings were predominant *bottom-up*. These patterns were largely preserved across coupling types (PAC and AAC) and across stimulus types (natural and synthetic auditory stimuli), suggesting they are a general feature of information processing in auditory cortex. Moreover, our findings showed that low-frequency PAC alternated between predominantly top-down or bottom-up over time. Altogether, this suggests sensory information need not be propagated along separate frequencies upwards and downwards. Rather, information can be unmixed by having low frequencies couple to distinct frequency ranges in the target region, and by alternating top-down and bottom-up processing over time.

**Significance:** The brain consists of highly interconnected cortical areas, yet the patterns in directional cortical communication are not fully understood, in particular with regards to interactions between different signal components across frequencies. We employed a a unified, computationally advantageous Granger-causal framework to examine bi-directional cross-frequency interactions across four sectors of the auditory cortical hierarchy in macaques. Our findings extend the view of cross-frequency interactions in auditory cortex, suggesting they also play a prominent role in top-down processing. Our findings also suggest information need not be propagated along separate channels up and down the cortical hierarchy, with important implications for theories of information processing in the brain such as predictive coding.

## 2 Introduction

One fundamental yet poorly understood component of cortical computation is the presence of widespread reciprocal cortical connections [Scott et al., 2015, Romanski and Averbeck, 2009, Kveraga et al., 2007, Salin and Bullier, 1995]. Moreover, most previous work has focused on the visual pathways. It remains unknown whether there are fundamental differences between top-down and bottom-up information processing along the auditory pathway. Cross-regional top-down effects in particular have remained unexplored, as most work has focused on bottom-up [Hyafil et al., 2015a, Giraud and Poeppel, 2012] or intra-regional effects [Lakatos et al., 2016]. In the present study, we examined bi-directional information processing across the macaque auditory hierarchy through an analysis of cross-frequency coupling (CFC). We computed two types of CFC measures, amplitude-amplitude (AAC) and phase-amplitude (PAC) coupling, for both natural and synthetic stimuli.

Cross-frequency coupling refers to correlation between the phase or amplitude component of one frequency and the amplitude component of other frequencies. This differs from linear analyses like coherence, which only examine coupling between signals at a single frequency. CFC, in contrast, considers interactions across the frequency spectrum. CFC may occur in several forms [Jirsa and Mueller, 2013] and has been observed during various tasks across several species including humans, macaques, and rodents. PAC has received widespread attention. Coupling of low-frequency (e.g. *θ, α*) phase to high-frequency (e.g. *γ*) amplitude has been associated with various behavioral mechanisms including learning, memory and attention [Hyafil et al., 2015b, Colgin, 2015, Doesburg et al., 2012, Kendrick et al., 2011, Canolty and Knight, 2010, Tort et al., 2009, Tort et al.,2008, Cohen, 2008, Jensen and Colgin, 2007, Canolty et al., 2006, Schack et al., 2002]. The reason for interest in PAC is that it may offer a mechanistic account of information coordination across neural populations and timescales. The phase of low-frequency oscillations could modulate the amplitude of high-frequency oscillations, putatively allowing control of spiking activity [Buzsáki et al., 2012, Ray and Maunsell, 2011]. AAC has also been shown to correlate with behavior [Canolty and Knight, 2010], but has been less explored mechanistically. Here we obtained estimates of both PAC and AAC strength as measures of cross-regional information processing in the bottom-up and top-down direction, allowing the two measures to be contrasted and compared in terms of informational content.

One prevalent theory of information processing in the brain which separates bottom-up and top-down components is *predictive coding*. In this (Bayesian) view of the brain, expectations are formed about incoming sensory information. These top-down predictions (expectations) are compared with bottom-up sensory information (outcomes) and representations are then potentially updated based on a prediction error (surprise), which is the difference between sensory inputs and their expectations. Neural implementations of predictive coding suggest that lower-order feedforward prediction errors are carried by a high frequency (e.g. *γ*) rhythm, while higher-order areas incorporate these errors into predictions which are fed back by a relatively lower frequency rhythm (e.g. *θ,α,β*) [Bastos et al., 2012, Friston, 2008]. Several studies have supported this view [Michalareas et al., 2016, Bastos et al., 2015, Bastos et al., 2012, Fontolan et al., 2014], though not without exceptions.

Here we probe this prediction by examining differences in cross-frequency coupling strength in the bottom-up and top-down direction across the frequency spectrum. It remains unknown whether this prediction can be generalized to cross-frequency interactions. Based on the predictive coding framework [Bastos et al., 2012, Friston, 2008], we would expect the phase of the modulating (source) rhythm to be *lower* in frequency (e.g. *θ,α,β*) in the top-down direction, and *higher* in frequency (e.g. *γ*) in the bottom-up direction. Alternatively, the same frequency bands could be employed in both directions. This may be possible by coupling to different ranges of the target amplitude spectrum, for instance. Also, bottom-up and top-down coupling could alternate in time, as has been suggested in a previous study [Fontolan et al., 2014].

Previous work has shown differences in how auditory cortical areas process natural and synthetic sounds [Anonymous, 2014]. It is possible that these differences translate into systematic differences in CFC patterns. If the revealed CFC pattern is a more general hallmark of inter-areal communication though, it should not be specific to stimulus type: natural and synthetic stimuli should show similar overall coupling patterns.

Our work revealed signature patterns of top-down and bottom-up processing in cross-regional PACs and AACs. These patterns were conserved across coupling types and across natural and synthetic stimuli, although overall reduced coupling strength was found for synthetic stimuli. We found that low frequencies featured prominently in both top-down and bottom-up information processing across the auditory hierarchy.

## 3 Methods and Materials

### 3.1 Subjects

Three adult male rhesus monkeys (Macaca mulatta) weighing 5.5–10 kg were used for recordings. All procedures and animal care were conducted in accordance with the Institute of Laboratory Animal Resources Guide for the Care and Use of Laboratory Animals. All experimental procedures were approved by the National Institute of Mental Health Animal Care and Use Committee.

### 3.2 Stimuli

The stimuli used for the main experiment included 20 conspecific monkey vocalizations (VOC) and two sets of 20 synthetic stimuli each (envelope-preserved sound, EPS; and spectrum preserved sound, SPS) derived from the original VOC stimuli. The VOC stimulus set consisted of 20 macaque vocalizations employed in a previous study [Kikuchi et al., 2010].

In order to obtain EPS from VOC stimuli, the envelope of a particular vocalization was estimated based on the amplitude component of the original stimulus’ Hilbert transform. The amplitude envelope was then multiplied by broadband white noise to create the EPS stimulus. Thus, all 20 EPS stimuli exhibited flat spectral content; these stimuli could not be discriminated based on spectral features, while the temporal envelopes (and thus the durations) of the original vocalizations were preserved.

SPS stimuli were obtained by first generating broadband white noise with a duration of 500 ms and computing its Fourier transform. The SPS stimulus’ amplitude in the Fourier domain was then replaced by the average amplitude of the corresponding VOC stimulus, before transforming back to the time domain by computing the inverse-Fourier transform. This resulted in a sound waveform that preserved the average spectrum of the original vocalization, while exhibiting a flat temporal envelope, random phase, with a duration of 500 ms. A 2 ms cosine rise/fall was then imposed on the stimulus to avoid abrupt onset/offset-effects. Hence, all 20 SPS stimuli exhibited nearly identical, flat temporal envelopes; these stimuli could not be discriminated using temporal features, while the average spectral power of the original vocalizations was preserved.

A total of 60 different stimuli were presented in pseudorandom order, with an interstimulus interval of 3 s. Each stimulus was presented 60 times. The sound pressure levels of the stimuli measured by a sound level meter ranged from 65 to 72 dB at a location close to the animal’s ear. Stimulus duration ranged from around .15 to 1 second (Anonymous, 2014).

### 3.3 Recordings

Custom-designed *µ*ECoG arrays (NeuroNexus Technologies, MI, USA) were used to record field potentials from macaque auditory cortex. Arrays were machine fabricated on a very thin polyimide film (20 *µ*m), with each array featuring 32 recording sites, 50 *µ*m in diameter each, on a 4 × 8 grid with 1 mm spacing (i.e., 3 × 7 mm rectangular grid) (Anonymous, 2014, 2012). Two animals, monkeys B and K, were implanted with arrays each in the left hemisphere, while one animal, monkey M, was implanted with arrays in the right hemisphere. Three of the arrays in each monkey were placed on top of STP, in a caudorostrally oriented row (Anonymous, 2014). The implantation procedure was described in detail previously (Anonymous, 2014, 2012).

### 3.4 Stimulus presentation and recording parameters

The experimental procedure was described in detail previously [Anonymous, 2014]. The monkey was placed in a sound-attenuating booth for the experiment (Biocoustics Instruments). The sound stimuli were presented while the monkey sat in a primate chair and listened passively with its head fixed. Auditory evoked potentials from the 128 channels of the ECoG-array were bandpassed between 2-500 Hz, digitally sampled at 1500 Hz, and stored on hard-disk drives by a PZ2–128 preamplifier and the RZ2 base station (Tucker Davis Technology).

### 3.5 Data Pre-Processing

Data analysis was performed using MATLAB R2017a (MathWorks) software. Since there was little significant auditory evoked power above 250 Hz, recordings were low-pass filtered and resampled at 500 Hz to enhance calculation speed and reduce memory requirements. The signal was bandpassed at 60Hz with a narrow-band filter to eliminate the possible presence of line-noise.

The 96 sites on the STP were grouped based on the characteristic frequency maps obtained from the high-gamma power of the evoked response to a set of pure-tone stimuli [Anonymous,2012,2014]. The change in frequency tuning along the STP reverses across areal boundaries, and these reversals were therefore used to identify the areal boundaries. This resulted in a grouping of electrodes into four sectors (Fig 1), which were estimated to correspond to the following subdivisions at a caudorostral-level within macaque auditory cortex: Sec (Sector) 1, A1/ML (primary auditory cortex/ middle lateral belt); Sec 2, R (rostral core region of the auditory cortex)/AL (anterior lateral belt region of the auditory cortex); Sec 3, RTL (lateral rostrotemporal belt region of the auditory cortex); Sec 4, RTp (rostro-temporal pole area). The recorded signal from each site partitioned into sectors 1-4 was re-referenced by subtracting the average of all sites within a particular sector [Fontolan et al., 2014, Kellis et al., 2010].

**Fig 1.**
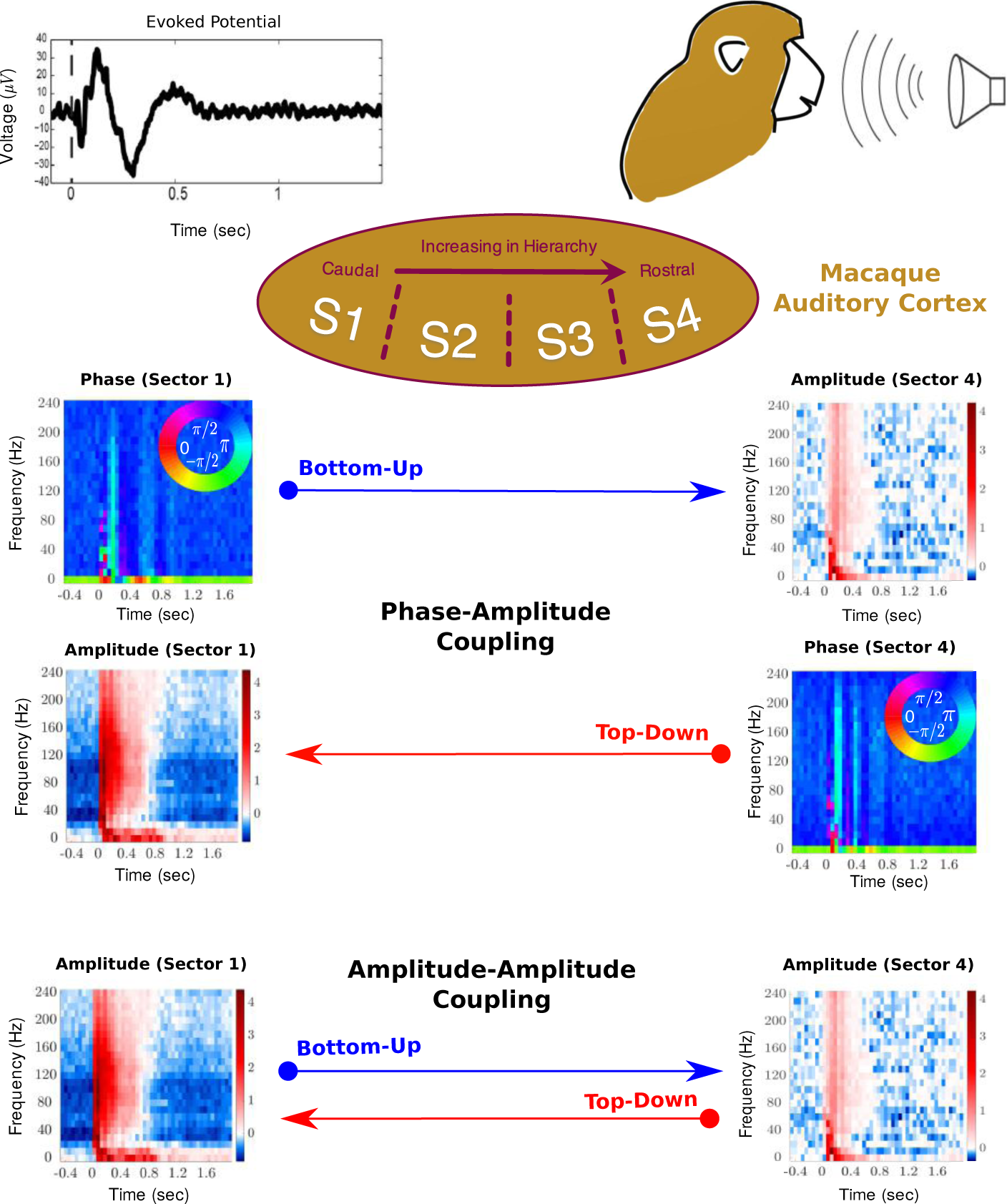
Coupling Analysis Overview: Recordings were made from four *µ*ECoG arrays spanning auditory cortex while monkeys listened to one of 20 natural vocalizations (VOC), 20 synthetic envelope-preserved sounds (EPSs) or 20 synthetic spectrum-preserved sounds (SPSs). Electrodes were partitioned into four sectors along the caudorostral axis (S1: A1/ML (primary auditory cortex/middle lateral belt), S2: R (rostral core region of the auditory cortex / AL (anterior lateral belt region of the auditory cortex), S3: RTL (lateral rostrotemporal belt region of the auditory cortex), S4: RTp (rostrotemporal pole area)). We decomposed the signal into its time-frequency representation, and obtained amplitude and phase components for each sector. We investigated two types of coupling across sectors: amplitude-amplitude and phase-amplitude coupling. We computed both types of coupling in the bottom-up and top-down direction. Top-down coupling was defined as coupling in which the source electrode came from a sector of higher order than the target electrode, and vice versa for bottom-up coupling.

We defined the following terminology to refer to particular frequency spaces throughout this paper: *delta* range: 0 - 2.5 Hz; *theta* range: 2.5 - 7.5 Hz; *alpha* range: 7.5 - 12.5 Hz; *low beta* range: 12.5 - 22.5 Hz; *high beta* range: 22.5 - 42.5 Hz; *low gamma* range: 42.5 - 67.5 Hz; *high gamma* range: *>* 67.5 Hz. The delta frequency band (0 - 2.5 Hz) was not included in the analysis since the pre-amplifier employed during the recording sessions does not allow for recording frequency components below 2 Hz and the power in this band was expectedly low.

### 3.6 Phase-amplitude (PAC) and amplitude-amplitude (AAC) cross-frequency coupling (CFC) Calculation

#### 3.6.1 Frequency Transformation

For Figures 1-5, the field potential data was first transformed into frequency space by computing the fast Fourier transform (FFT) of the signal in every channel using a 200 ms Hanning window, stepped by 200 ms. Thus, we obtained a frequency representation of the data at a 5Hz bandwidth ranging from 2.5Hz - 132.5Hz, which satisfies the Nyquist-criterium. We obtained the phase and amplitude information in a given source and target region for every channel and normalized the amplitude information in each frequency band by the corresponding power in that frequency band, averaged across the entire experiment, so as to account for decreasing power with frequency (Fig. S1).

#### 3.6.2 Obtaining CFCs using canonical correlation

We computed two types of cross-frequency coupling, phase-amplitude (PAC) and amplitude-amplitude (AAC) coupling. Couplings were calculated for every channel pair across the four sectors (S1-S4), resulting in 6 possible cross-sector pairs (S1-S2, S1-S3, S1-S4, S2-S3, S2-S4, S3-S4). We obtained estimates of coupling strength in two directions, bottom-up and top-down. In the former case, the source component was derived from a recording site in a sector lower in the auditory hierarchy than the target component (referred to as bottom-up coupling), while in the latter case the source component came from a sector higher in the auditory hierarchy than the target component (referred to as top-down coupling) (Fig. 1). Ultimately, we were interested in examining the difference in coupling strength in the top-down and bottom-up directions across the auditory hierarchy. Results are presented collapsed across all cross-sector pairs (Fig. 2), as well as separately for every cross-sector pair (Fig. 3), averaged across all channel pairs and across the three animals in both cases.

**Fig 2.**
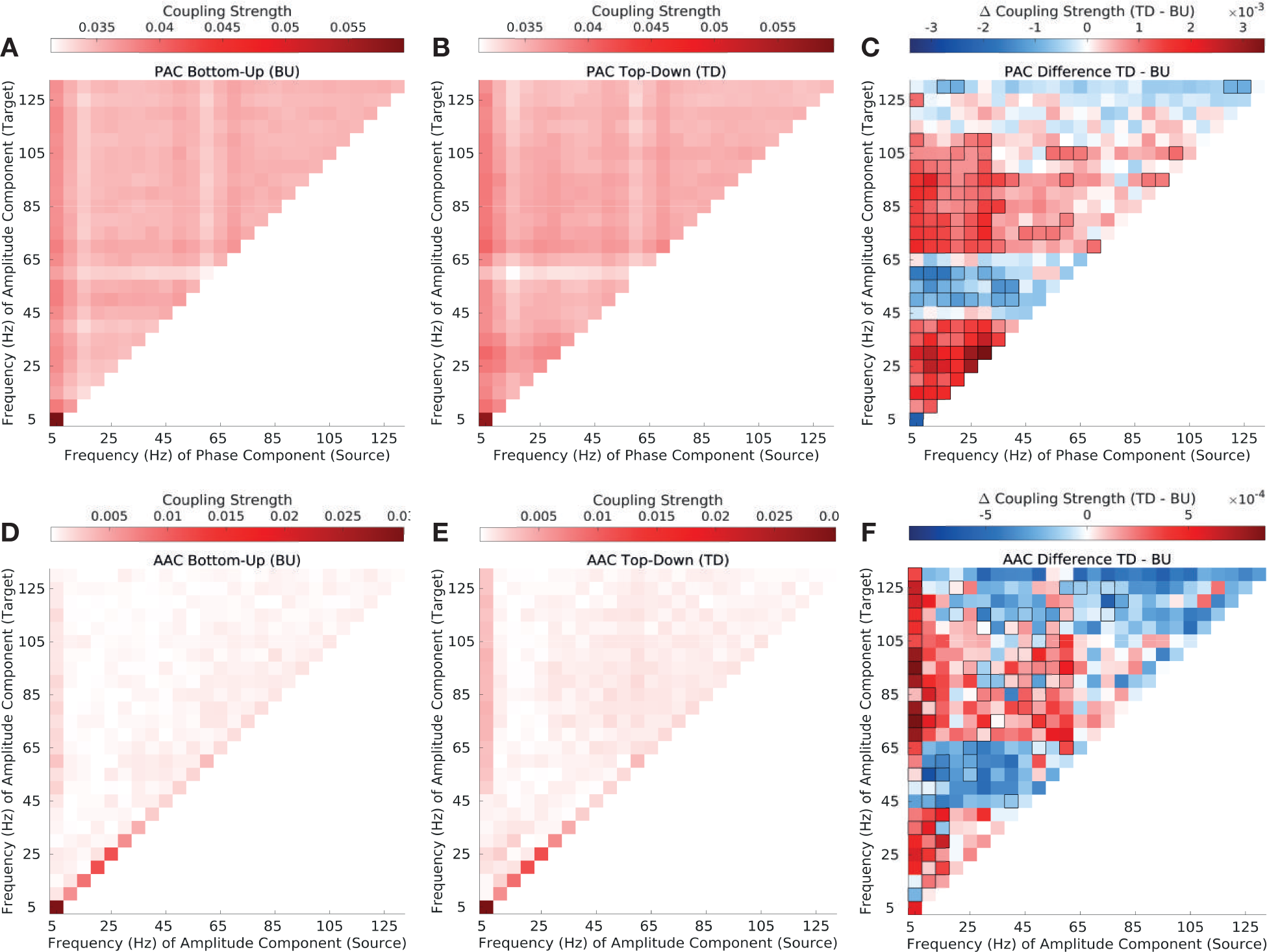
Top-down vs. bottom-up phase-amplitude and amplitude-amplitude coupling in natural vocalizations (VOC). **(A-B)** Phase-amplitude coupling (PAC) strength in the top-down (A) and bottom-up direction (B). Depicted are canonical correlation-derived coupling coefficients (see Methods and Materials). **(C)** Difference in top-down and bottom-up in PAC strength. Significant differences are enclosed by black rectangles (FDR corrected, q-value=0.01). **(D-E)** Amplitude-amplitude coupling (AAC) strength in the top-down (D) and bottom-up direction (E). Depicted are canonical correlation-derived coupling coefficients. **(F)** Difference between top-down and bottom-up in AAC strength. Significant differences are enclosed by black rectangles (FDR corrected, q-value=0.01). Results are depicted averaged across all channels, cross-regional pairs, and animals.

**Fig 3.**
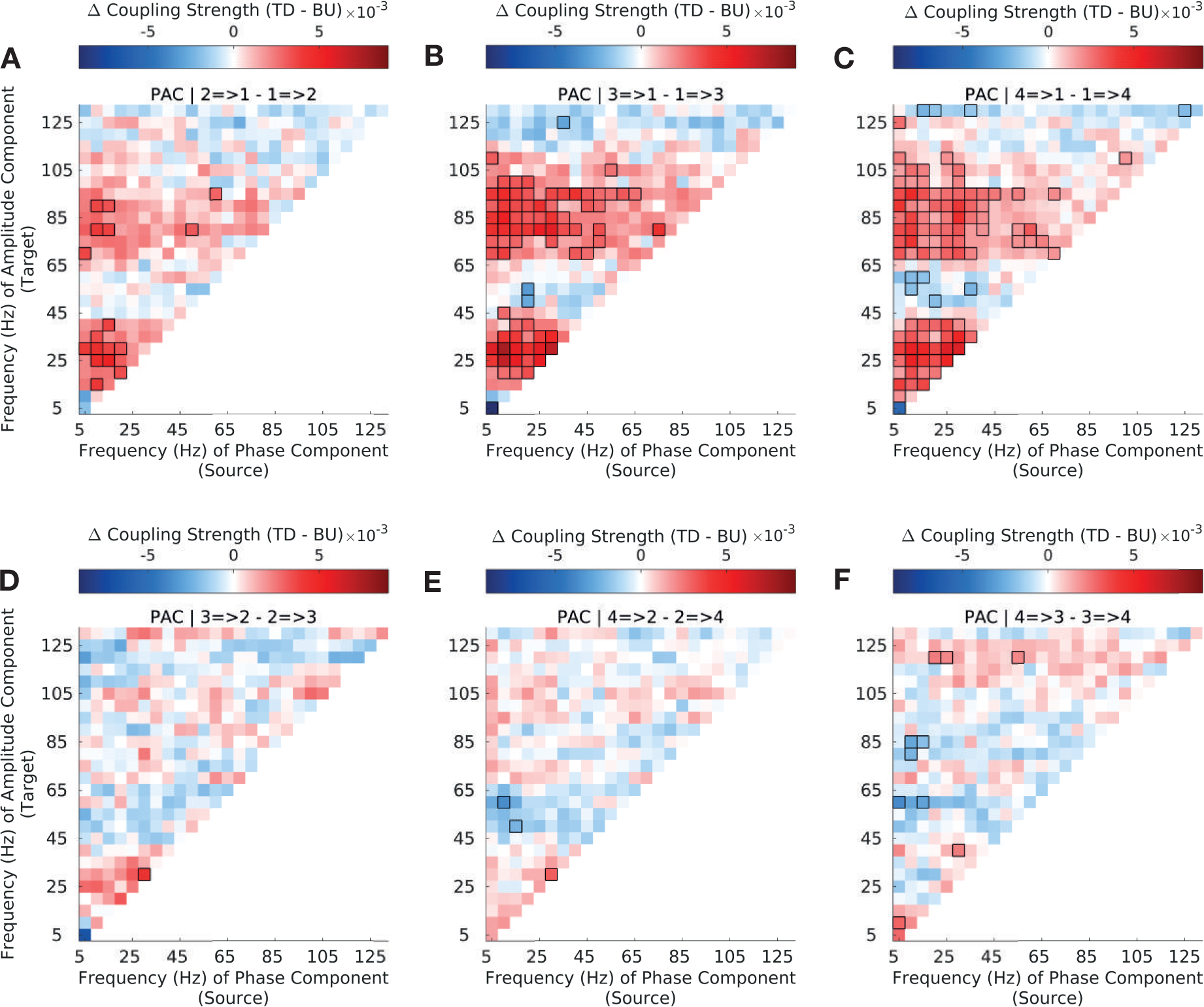
Top-down versus bottom-up phase-amplitude coupling strength (PAC) across the auditory hierarchy in natural stimuli. Difference in top-down and bottom-up *phase-amplitude* coupling (PAC) strength for CFC between sectors 1 (A1/ML) and 2 (R/AL) **(A)**, sectors 1 (A1/ML) and 3 (RTL) **(B)**, sectors 1 (A1/ML) and 4 (RTp) **(C)**, sectors 2 and 3 **(D)**, sectors 2 and 4 **(E)** and sectors 3 and 4 **(F)** (see Methods and Materials 3.5 for sector definitions). Interactions between sector 1 (A1/ML) and higher order sectors show strong asymmetries in bottom-up and top-down coupling strength across the frequency spectrum; interactions among higher-order sectors (S2-S4) show less widespread asymmetries. Significant differences are enclosed by black rectangles (FDR corrected, q-value=0.01). Results are depicted averaged across all channels and animals.

We used canonical correlation analysis (CCA), to compute both types of coupling. The advantage of CCA is that it computes coupling among multivarite observations in the source and target regions. This framework, therefore, allows CFC values to be computed across the entire frequency range simultaneously. Furthermore, some approaches to calculating AAC and PAC first carry out dimensionality reduction on the source and target region signals separately, and then calculate correlations in the low dimensional space. If the variance preserving dimensions in the source and target space are not the dimensions that maximize correlations between the source and target space, interactions may be missed. CCA on the other hand finds the low dimensional representations that maximize coupling between the source (*X*) and target (*Y)* space. In CCA, one is seeking projections (linear combinations) *U* = *XA* and *V* = *Y B* such that the correlation *corr*(*U, V)* among these derived directions is maximal [Durstewitz, 2017].

We carried out our analyses using a Granger Causal framework. Therefore, we first carried out CCA between amplitude in the target region and time lagged versions of the amplitude (Eqs 1-2), obtaining an estimate of the the coupling strength *Ŷ* that could be inferred purely from coupling of the signal at the target electrode onto itself. We used the first singular value only to obtain *Ŷ*; results did not differ from those obtained from including all entries. We then obtained the residual from this analysis 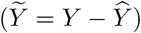, and used this residual when we examined the interactions between regions. Subsequent analysis was performed using the residuals Ŷ from this regression as the signal for the target region.

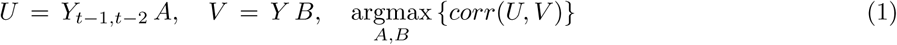

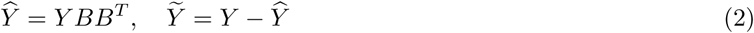

We then computed phase-amplitude (PAC) and amplitude-amplitude (AAC) cross-frequency couplings across the four sectors (Fig. 1), by regressing the phase or amplitude component of the signal from the source sector, respectively, on the amplitude component of the signal in the target sector for every channel. PAC and AAC was calculated in the CCA framework across all time windows and trials altogether, simultaneously across all frequency bands. In the case of PAC, *X* contained the sine and cosine transformed phase component of the signal, while *Y* contained the squared and log-transformed amplitude component of the signal, with *n* = 10800 (1200 trials * 9 windows) observations for VOC and SPS stimuli and *n* = 10773 observations (1197 trials * 9 windows) for EPS stimuli, and *k* = 25 *\** 2 and *k* = 25 predictors for PAC and AAC, respectively (Eq 3).

After obtaining the canonical coefficient matrices *B* and *A* within the CCA framework (Eq 3), one can obtain the coupling matrix *P* by multiplying the *B* and *A* matrices with the singular values *S* of the correlation matrix (Eq 4). We used the first 10 directions (and singular values of *S*) that captured most of the interaction; results did not differ from those obtained from including all entries. For AAC, the final cross-frequency coupling matrix is *P*. For PAC, one combines the sine and cosine components of *P* by computing their Euclidian norm, yielding the final coupling matrix 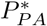.

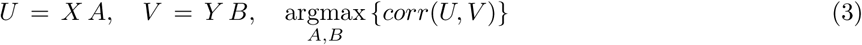

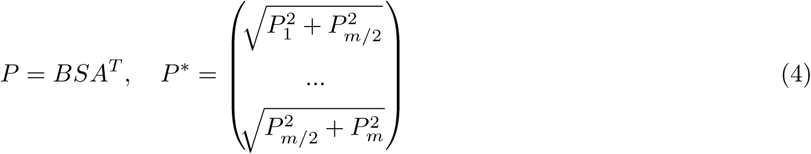

#### 3.6.3 Obtaining the difference between top-down (TD) and bottom-up (BU) coupling

We obtained AAC and PAC coupling matrices in this manner for both the bottom-up and the top-down direction. In bottom-up coupling, the source component was derived from a recording site in a sector lower in hierarchy than the target component (referred to as bottom-up coupling), while in top-down coupling, the source component came from a sector higher in hierarchy than the target component (Fig. 1). We then obtained the difference Δ*P*_*same* *stim*_ between the top-down and bottom-up coupling matrices *P*_*Top-Down*_ and *P*_*Bottom-Up*_ (Eq 5). This was done separately for each of the cross-sector pairs. Results are presented collapsed across all cross-sector pairs (Fig. 2), as well as separately for every cross-sector pair (Fig. 3).

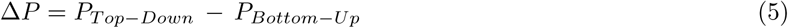

#### 3.6.4 Obtaining the difference in coupling strength between original (VOC) and synthetic stimuli (EPS/SPS)

In order to examine the difference in coupling strength between natural and synthetic stimulus types (Fig. 4,5), we computed the difference between VOC and EPS stimuli (Fig. 4) and between VOC and SPS stimuli (Fig. 5) separately in the top-down (TD, Eq 6) and bottom-up direction (BU, Eq 7).

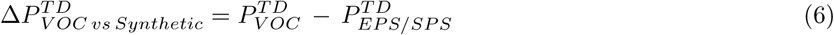

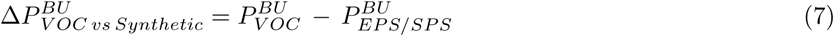

**Fig 4.**
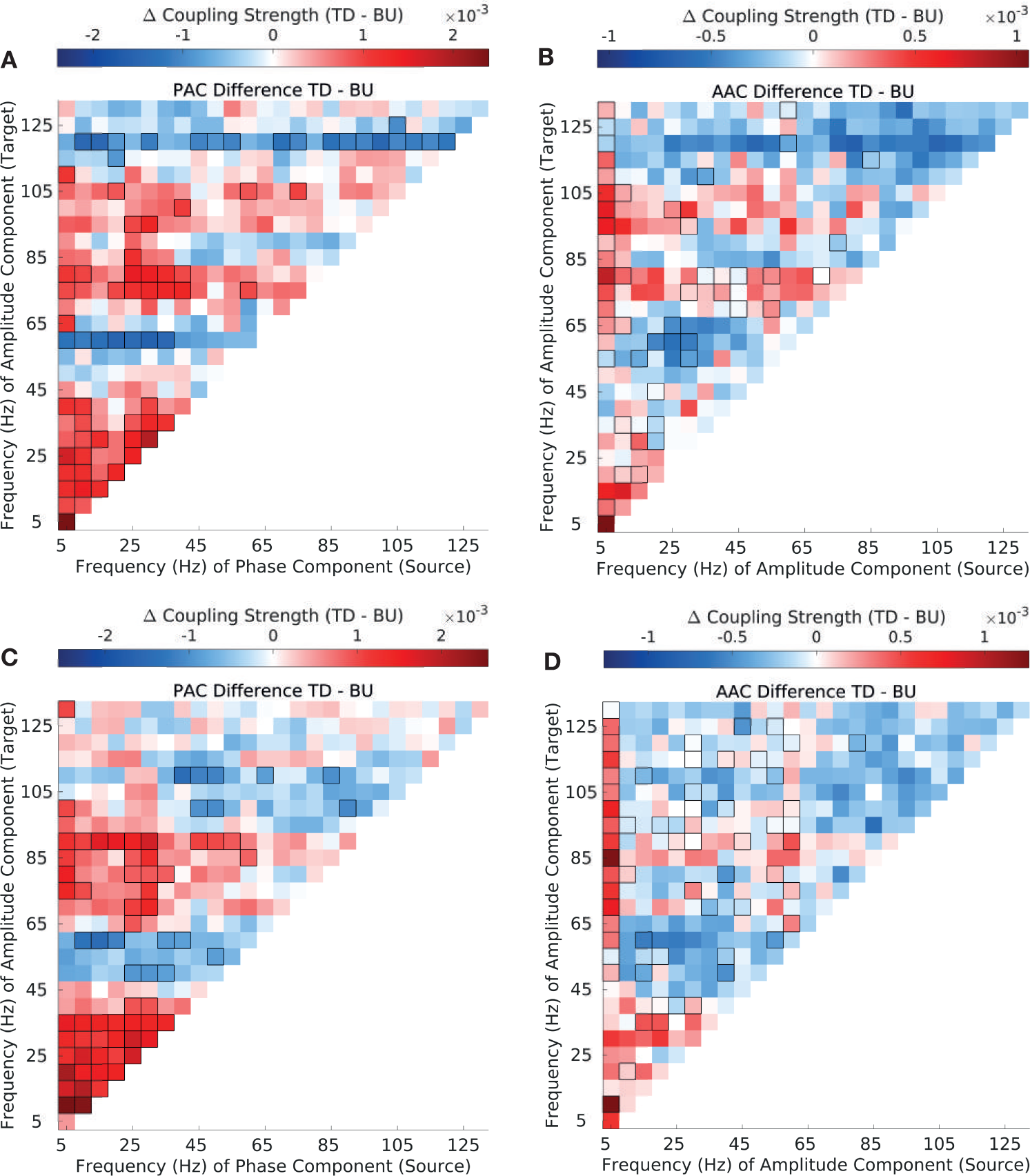
Top-down vs. bottom-up phase-amplitude and amplitude-amplitude coupling in synthetic envelope-preserved stimuli (EPS) and synthetic spectrum-preserved stimuli (SPS) **(A)** Difference in top-down and bottom-up PAC strength in EPS stimuli. **(B)** Difference in top-down and bottom-up AAC strength in EPS stimuli. **(C)** Difference in top-down and bottom-up PAC strength in SPS stimuli. **(D)** Difference in top-down and bottom-up AAC strength in SPS stimuli. Significant differences are enclosed by black rectangles (FDR corrected, q-value=0.01). Results are depicted averaged across all channels, cross-regional pairs, and animals.

**Fig 5.**
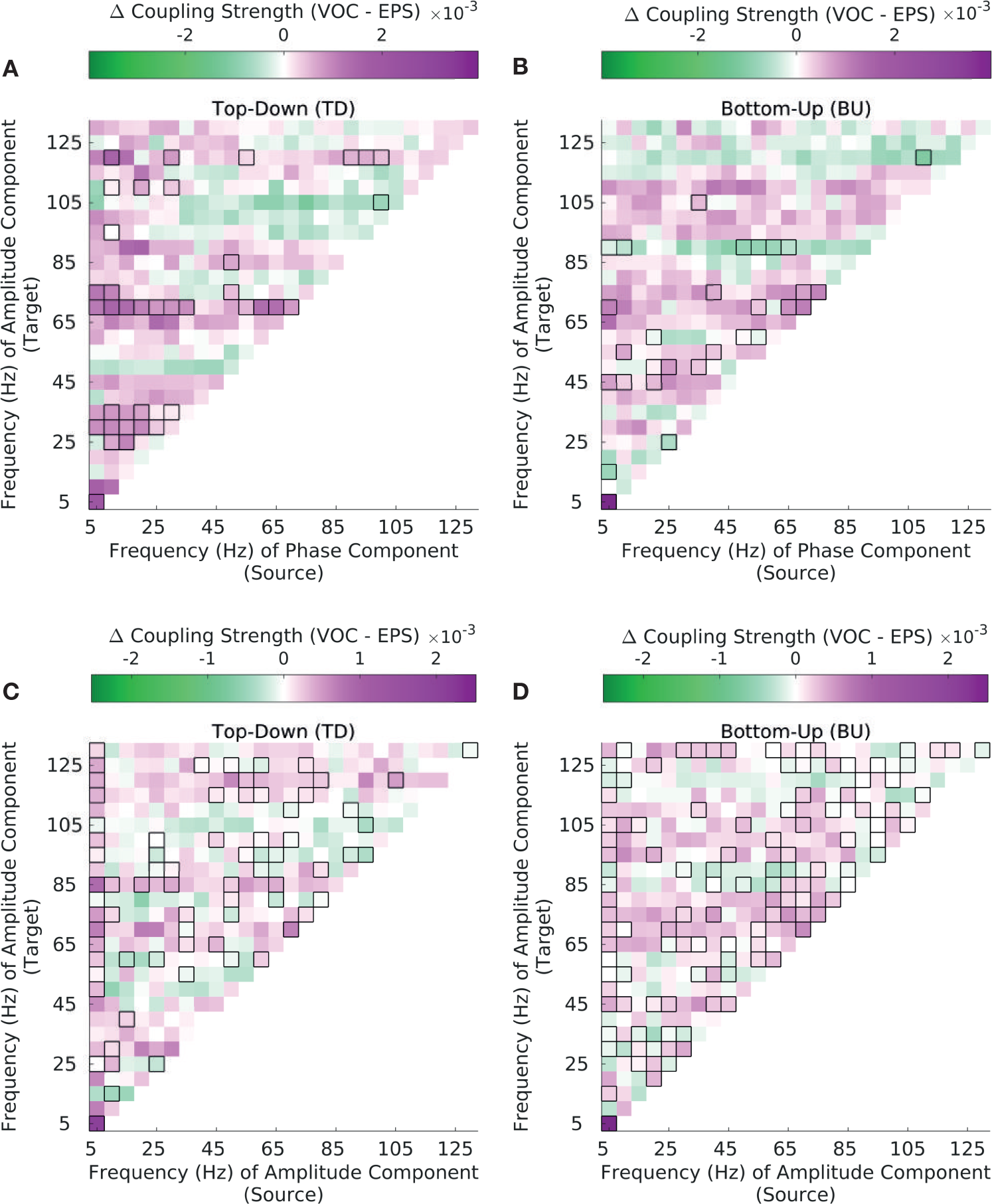
Top-down versus bottom-up phase-amplitude and amplitude-amplitude coupling strength in synthetic envelope-preserved sounds (EPS). **(A-B)** Difference in PAC strength between natural (VOC) and synthetic envelope-preserved sounds (EPS), separately in the top-down (B) and bottom-up direction (C). **(C-D)** Difference in AAC strength between VOC and EPS, separately in the top-down (C) and bottom-up direction (D). Significant differences are enclosed by black rectangles (FDR corrected, q-value=0.01). Results are depicted averaged across all channels, cross-regional pairs, and animals.

### 3.7 PAC cross-frequency coupling over time

#### 3.7.1 Frequency Transformation

For computing cross-frequency coupling over time (Fig. 6), the field potential data was first transformed into frequency space by bandpassing the signal and subsequently employing the Hilbert transform. Here, we wanted to examine modulation of the amplitude signal in the target region by a *θ* (2.5Hz-7.5Hz) phase rhythm from the source region (cross-sector PAC); we focused on the *θ* band since it contained the most power (Fig. S1). With this in mind, we bandpassed the signal from the source region at a 5Hz bandwidth and the signal from the target region at a 10Hz bandwidth with zero-phase digital filtering (Matlab filtfilt). The bandwidth of the amplitude component in the target region was chosen to be twice as wide as the bandwidth of the modulating phase component in order to ensure the amplitude bandwidth includes the side peaks produced by modulating *θ*-range phase rhythm [Aru et al., 2015]. The signal from both source and target regions was subsequently transformed into the frequency space by applying the Hilbert transform. We then obtained the instantaneous phase and amplitude signal for a given source and target region, respectively, in every channel. It is possible that due to the nature of the filter choice, signal leaked to earlier time windows (in particular to before stimulus onset) thus leading to higher coupling estimates; however, we were interested in examining differences in bottom-up and top-down coupling here, and signal leak would have affected both directions equally, in other words there is no a priori reason to expect coupling estimates to be higher in one direction over the other.

**Fig 6.**
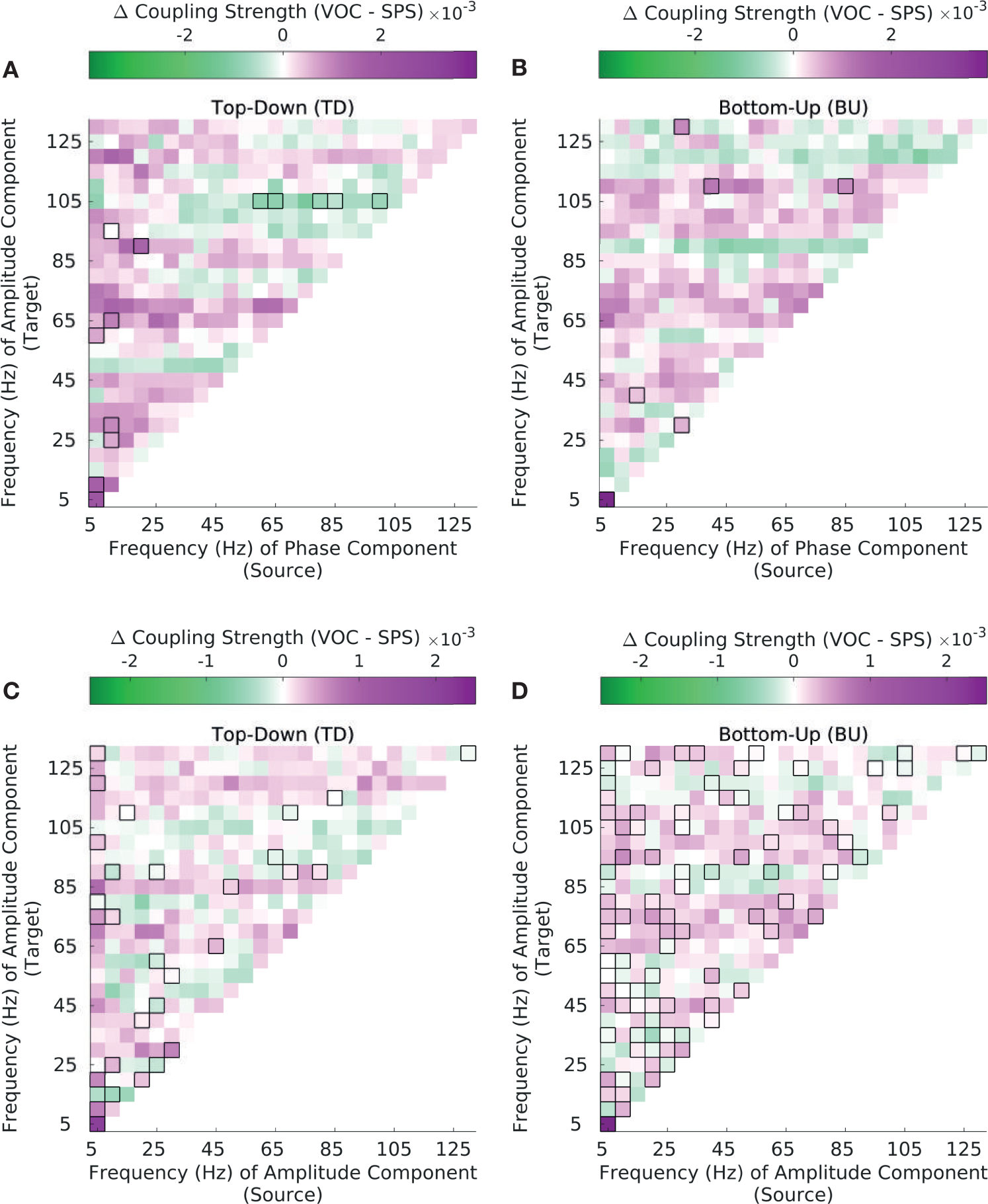
Top-down versus bottom-up phase-amplitude and amplitude-amplitude coupling strength in synthetic spectrum-preserved sounds (SPS). **(A-B)** Difference in PAC strength between natural (VOC) and synthetic spectrum-preserved sounds (SPS), separately in the top-down (B) and bottom-up direction (C). **(C-D)** Difference in AAC strength between VOC and SPS, separately in the top-down (C) and bottom-up direction (D). Significant differences are enclosed by black rectangles (FDR corrected, q-value=0.01). Results are depicted averaged across all channels, cross-regional pairs, and animals.

As a check to ensure our results were not driven by filter choice, we also obtained FFT-ed data at a 25Hz bandwidth ranging from 2.5Hz to 132.5Hz, using a window size of 40ms. The larger bandwidth size resulted from the necessity to achieve sufficient temporal resolution.

#### 3.7.2 Obtaining CFCs over time

CFCs were then computed within the canonical correlation framework as described above. PACs were estimated from the Hilbert-transformed (or FFT-ed) signal by computing the canonical correlation between the phase signal from the source region and the residual (from regressing out *Y*’s past as described above) of the amplitude signal from the target region. In contast to the analyses described above, now separate estimates for bi-directional coupling coefficients were obtained for every time window, using *n* = 1200 trials for VOC and *n* = 1197 trials for EPS/SPS.

### 3.8 Statistical Analysis

Paired t-tests were performed to establish statistical significance of the difference between top-down and bottom-up coupling (Δ*P*; Fig. 2C/F), as well as the differences between VOC and synthetic stimuli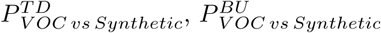; Fig.4,5), and the difference between bottom-up and top-down PAC across time (Fig. 6). This was done separately for each frequency pair, by testing whether the mean of the difference between top-down and bottom-up coupling was significantly different from zero (null hypothesis). Lilliefors’ test for normality was conducted for each frequency pair in order to ensure the normality assumption was satisfied, and histogram plots were inspected for select frequency pairs (Fig. S2,S3). In order to assess overall differences in coupling strength for natural and synthetic stimuli (Fig. 2C/F,Fig. 4,S4,S6,S7,S10,S11), paired t-tests were conducted across all channel pairs, cross-regional pairs and animals altogether. In order to assess coupling differences per regional pair (Fig. 3,S5,S8,S9,S12,S13), paired t-tests were conducted separately for each regional pair, across all channel pairs and animals altogether. When assessing coupling differences across time (Fig.6,S14), paired t-tests were conducted separately for every window, across all channel pairs and animals. Couplings significant post FDR-correction with *α* = 0.01 are highlighted with black contours in every figure. Additionally, as a more conservative test against type I errors, significance of the difference between top-down and bottom-up coupling (Δ*P)* was also computed using the cluster-permutation method (Fig. S4,S7,S11) [Maris and Oostenveld, 2007].

## 4. Results

We analyzed local field potentials recorded simultaneously from multiple cortical areas in the auditory cortex of three macaques while the animals listened to auditory stimuli (Fig. 1). These auditory stimuli consisted of 20 natural conspecific vocalizations, 20 envelope-preserved stimuli (EPS) and 20 spectrum-preserved stimuli (SPS), which were derived from the original vocalizations [Anonymous,2014]. A previous study established the approximate rostrocaudal location of implanted micro-electrocorticography (*µ*ECoG) arrays along the auditory hierarchy by the characteristic frequency of each electrode contact (Anonymous, 2014, 2012). This partitioned the recording sites into four sectors (S1-S4), putatively spanning the caudorostral levels from the core A1 (S1) to RTp (S4), including some of the surrounding belt areas (Fig. 1). We operationally defined feedforward to occur from earlier to later sectors (bottom-up direction), and feedback to occur from later to earlier sectors (top-down direction).

We wanted to examine cross frequency interactions across the auditory hierarchy. Phase-amplitude coupling is the correlation between the phase at a single frequency in one site, and the amplitude in another site, while amplitude-amplitude coupling is the correlation between the amplitude at a single frequency in one site, and the amplitude at another (and also the same) frequency at another site. When all frequency-frequeny interactions are considered, a matrix of coupling coefficients is generated. To carry out the analysis we first decomposed the signal from every trial and every site into its spectral components using the short-time fast Fourier transform. This resulted in time-phase and time-amplitude data (Fig. 1). Next, we obtained measures of phase-amplitude and amplitude-amplitude coupling strength using canonical correlation analysis (CCA), which allowed us to estimate coupling between all frequencies in the source area and all frequencies in the target area, simultaneously. Analyses were further carried out in a Granger Causal framework. Therefore, for all analyses we first estimated activity in the target region, using time lagged activity from the same region. We then calculated the residual from this analysis, and used these residuals when examining coupling from other areas (see Methods and Materials). Thus, the coupling coefficients represent correlations over and above coupling that could be inferred from past activity in the target region, on itself.

We computed coupling strength across the frequency spectrum for each cross-regional pair, separately in the top-down and bottom-up direction (Fig. 1). Top-down coupling was defined as coupling in which the source electrode came from a sector of higher order than the target electrode, and vice versa for bottom-up coupling. For example, coupling between the phase (or amplitude) in S4 and the amplitude in S1 is defined as top-down, and coupling between the phase (or amplitude) in S1 and the amplitude in S4 is defined as bottom-up in the case of PAC (or AAC). This definition is in accordance with a previous approach [Fontolan et al., 2014].

### 4.1 Distinct cross-frequency coupling signatures for bottom-up and top-down coupling, both involving low frequencies

We first examined PAC and AAC across the four sectors, in auditory evoked potentials to 20 conspecific vocalizations (VOC). Both CFC measures peaked in the *θ* frequency range and showed relatively strong coupling across all the target amplitude frequencies (Fig. 2A,B,D,E). The field potential signal showed the highest power in the same frequency band decaying with the typical 1*/f* pattern thereafter (Fig S1), thus satisfying a pre-requisite for the presence of physiologically meaningful CFC [Aru et al., 2015].

Among PACs, both top-down and bottom-up coupling was dominant between *θ* phase and broadband amplitude (Fig. 2A,B). Examining differences in coupling strength in the two directions more closely (Fig. 2C), PACs showed dominant top-down coupling in the low-frequency space (*θ, α, β* phase to *θ, α, β* amplitude), as well as between low frequency phase and high-*γ* (*>*67.5Hz) amplitude. Bottom-up PAC was dominant between low frequency phase and low-*γ* (47.5-62.5Hz) amplitude. The same trends were found when computing significance at p=0.01 using cluster-permutation testing (Fig. S4A). In addition, bottom-up coupling was dominant between *θ* phase and *θ* amplitude, low *β* (12.5-22.5) phase and high-*γ* amplitude (127.5-132.5Hz), and between high *γ* phase and high *γ* amplitude. These interactions did not show up significant with cluster-permutation testing (Fig. S4A) [Maris and Oostenveld, 2007].

AACs showed a similar pattern to PACs in both directions, but coupling strength and differences in coupling strength were overall lower than for PACs (Fig. 2D,E). AACs replicated the pattern oveserved in PACs with dominant *θ* to broadband coupling in both directions. Additionally, and unlike in PACs, coupling strength among same-frequency couplings was pronounced in both directions with strongest coupling in the low-frequency range (Fig. 2D,E). Examining differences in coupling strength in the two directions more closely (Fig. 2F), AACs displayed a similar pattern to PACs but showed less significant frequency pairs than PACs across the entire frequency spectrum. AAC coupling differences did not show up significant with cluster-permutation testing (Fig. S4B) [Maris and Oostenveld, 2007].

### 4.2 Cross-regional coupling mainly driven by interactions between A1 and higher order sectors

To examine cross-regional patterns in coupling across the auditory hierarchy (Fig. 1) more closely, we analyzed differences between top-down and bottom-up coupling strength separately across the six cross-regional pairs (Fig. 3). This revealed that the overall coupling pattern (Fig. 2) was mostly driven by interactions between S1 (A1/ML) and S2-S4 (higher-order sectors) (Fig. 3 A,B,C), rather than by interactions among higher-order sectors (Fig. 3 D,E,F).

More specifically, interactions between S1 (A1/ML) and S2 (R/AL) show dominant top-down coupling only (Fig. 3A), which is overall less widespread across low and high frequency amplitudes than for the cross-regional pairs S1 (A1/ML)-S3 (RTL) (Fig. 3B) and S1 (A1/ML)-S4 (RTp) (Fig. 3B). The full pattern as manifested in Fig. 1 only gains full prominence among cross-regional pairs S1-S3 (3B) and S1-S4 (3B), which also include predominant bottom-up coupling between low frequency phase and low-*γ* amplitude. Interactions between higher-order sectors (3 D,E,F) show fewer differences between top-down and bottom-up coupling. A similar trend was observed among AACs, though differences were overall smaller compared to PACs and there was less widespread low frequency to *highγ* coupling among S1-S3 and S1-S4 (Fig. S5).

### 4.3 Top-down and bottom-up cross-frequency signatures generalize to synthetic stimuli

To examine whether the observed coupling patterns are a general feature of inter-areal communication, we also analyzed coupling strength in two types of synthetic stimuli derived from the natural vocalizations. Envelope-preserved sounds (EPS) were obtained by leaving the original temporal envelope of the vocalization intact but flattening its spectral content, while spectrum-preserved sounds (SPS) were obtained by leaving the spectral information of the original vocalization intact but flattening its temporal envelope [Anonymous, 2014] (see Methods and Materials 3.2).

EPS and SPS stimuli showed similar bottom-up and top-down coupling patterns to natural stimuli, with pronounced *θ*-to-broadband coupling strength in both directions (Fig. S6,S10). Examining differences between top-down and bottom-up coupling strength in both types of synthetic stimuli (Fig. 4,S7, S11), we found that they reflected the same coupling asymmetries across the frequency spectrum as natural stimuli did (Fig. 2). This was true for PAC (Fig. 4A,C) and AAC (Fig. 4B,D) alike. More specifically, both types of synthetic stimuli showed dominant top-down coupling among low-to-low frequency and low-to-high *γ* interactions, and dominant bottom-up coupling among low-to-low *γ* interactions, like natural stimuli did. Both types of synthetic stimuli also showed similar coupling asymmetries to natural stimuli when examining cross-regional pairs separately (Fig. S8, S9, S12, S13), with most of the pattern driven by interactions between A1 and higher-order sectors, as in natural vocalizations.

### 4.4 Spectrally flat stimuli show more differences in bi-directional coupling to natural stimuli than spectrally intact stimuli

To examine differences in coupling strength between natural and synthetic stimuli more closely, we compared coupling strength across stimuli separately in the top-down and bottom-up direction (Fig. 5).

EPS stimuli showed lower coupling strength than natural stimuli in both directions for both PACs and AACs (Fig. 5). Among PACs more specifically (Fig. 5A-B), natural stimuli showed higher coupling strength in the top-down direction for select frequency pairs in the low frequency space as well as among low frequency to high-*γ* coupling (Fig. 5A). In the bottom-up direction, coupling strength for natural stimuli was enhanced among low frequency to low and high *γ* couplings (Fig. 5B). There were also select coupling values in the low and high *γ* frequency target amplitude space for which coupling was stronger in EPS stimuli.

Among AACs, coupling strength was increased for natural stimuli across the entire frequency spectrum in both directions (Fig. 5 C-D); compared to PACs, AACs showed more significant differences between natural and EPS stimuli in the high-frequency space, in both directions (Fig. 5 C,D). Altogether, coupling strength is enhanced for natural stimuli over envelope-preserved but spectrally flat synthetic stimuli (EPS) in both directions and across the entire frequency spectrum, with AACs displaying more significant differences in the high-frequency space.

SPS stimuli showed more similar coupling patterns to VOC stimuli. Among PACs, there were much less significant differences in both directions (Fig. 6A-B) than there were between EPS and natural stimuli (Fig. 5A-B). This also held true among AACs (Fig. 6C-D), especially in the top-down direction (Fig. 6C) which displayed much less significant differences across the frequency spectrum than for EPS stimuli (Fig. 5C). SPS and natural stimuli still showed differences among AACs in the bottom-up direction (Fig. 6D) across the entire frequency spectrum, mostly with increased coupling strength for natural stimuli. Altogether, spectrally rich (but envelope flat) stimuli (SPS) showed less widespread differences to natural stimuli in both directions for both types of coupling; most of the differences were observed in the bottom-up direction among AACs, with natural stimuli showing higher coupling strength across the frequency spectrum.

Overall, top-down and bottom-up cross-frequency coupling signatures were preserved among synthetic stimuli, albeit at overall reduced coupling strength. Envelope-flattened (but spectrally intact) SPS stimuli showed smaller differences in coupling strength to VOC stimuli than spectrally-flat (but envelope preserved) EPS stimuli. AACs revealed more widespread differences in coupling strength than PACs among synthetic stimuli, in particular in the *γ* frequency range.

### 4.5 Top-down and bottom-up coupling alternate across time

To examine the pattern of PAC coupling over time in more detail, we focused on the *θ* frequency phase band which showed the highest coupling strength (Fig. 2A-B). We bandpassed the signal and obtained instantaneous phase and amplitude information by applying the Hilbert-transform (as described in Methods and Materials 3.7). We then computed PAC for every time window separately, obtaining a time-resolved picture for coupling between the *θ* (2.5-7.5Hz) phase band in the source sector and *θ*/*α* (2.5Hz-12.5Hz) amplitude band in the target sector (Fig. 7).

**Fig 7.**
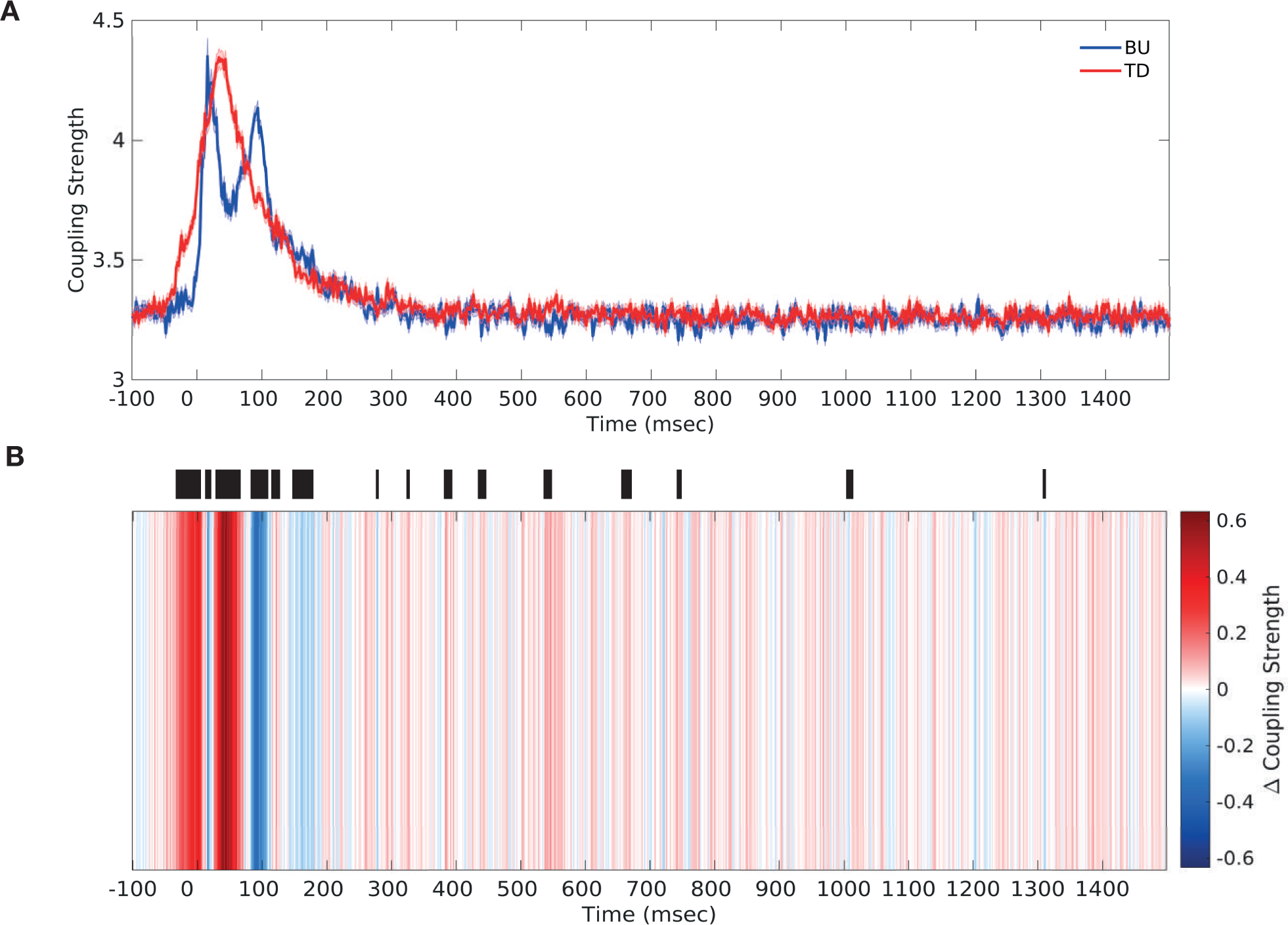
Top-down vs. bottom-up low-frequency PAC over time. **(A)** *θ* (2.5-7.5Hz) phase to *θ*/*α* (2.5-12.5Hz) amplitude PAC over the duration of stimulus presentation, separately in the bottom-up direction (blue) and the top-down direction (red) **(B)** Differences in top-down and bottom-up low frequency PAC (*θ* (2.5-7.5Hz) phase to *low γ* amplitude coupling) over the duration of stimulus presentation. Significant differences are marked by thick black lines (FDR corrected, q-value=0.01). Results are depicted averaged across all channels, cross-regional pairs, and animals. Top-down and bottom-up coupling were found to alternate in the period around stimulus onset (−50ms to 200ms).

Both bottom-up and top-down coupling strength (Fig. 7A) was increased in the period around stimulus onset (−50ms to 200ms). Examining the difference in coupling strength in the two directions over time (Fig. 7B), we observed that top-down coupling was enhanced immediately before and during stimulus onset (−50 to 10ms), followed by a short period of enhanced bottom-up coupling post stimulus onset. This was again followed by a bout of enhanced top-down coupling (around 50ms), with top-down coupling peaking while bottom-up coupling decreased (Fig. 7A). Thereupon, bottom-up coupling was enhanced again (around 100ms), with bottom-up coupling peaking while top-down coupling was decreasing (Fig. 7A).

When examining differences in coupling strength over time at the level of individual cross-regional pairs (Fig. S14), we found an alternating pattern across all but one pair though the exact pattern differed somewhat across pairs. The second peak of bottom-up coupling (around 100ms) was present for all pairs but for interactions between S1 and S2 (Fig. S14A). Hence, this second peak of bottom-up coupling could represent the delayed processing of bottom-up information as signal propagates to higher-order sectors.

We also computed coupling over time from Fourier-transformed, instead of Hilbert transformed, data (see Methods and Materials) to make sure the effect we were observing was not was not a by-product of filter choice, and we obtained a similar result (Fig. S15). Bi-directional coupling strength was enhanced in the period around stimulus onset, with enhanced top-down coupling before and during stimulus onset followed by enhanced bottom-up coupling around 100ms (Fig. S15A,B). Hilbert- and Fourier-transformed results are not directly comparable, however, due to the different time/frequency trade-offs for the two methods. Overall, we observed that top-down and bottom-up coupling alternated in dominance in the period around stimulus onset.

## 5. Discussion

We found distinct cross-frequency signatures for top-down and bottom-up information processing in auditory cortex. These patterns were similar across phase-amplitude and amplitude-amplitude couplings, and across stimulus types (albeit at lower coupling strength in synthetic spectrum-flat stimuli). More specifcally, we observed dominant top-down coupling among low-to-low and low-to-high *γ* frequency interactions, and dominant bottom-up coupling among low-to-low *γ* frequency interactions. Spectrally intact (but envelope-flattened) synthetic stimuli showed smaller differences in coupling strength to natural vocalization stimuli than did spectrally flat (but envelope preserved) synthetic stimuli. AACs showed more widespread differences in coupling strength than PACs among synthetic stimuli, in particular in the *γ* frequency range. Altogether, our results suggest these cross-frequency signatures are a general hallmark of bottom-up and top-down information processing in auditory cortex. Moreover, they suggest information need not be propagated along separate frequency bands up and down the auditory hierarchy; rather, low frequencies (*θ,α,β*) may be prominently involved in both bottom-up and top-down information processing by coupling to different ranges of the target frequency spectrum, and by alternating bottom-up and top-down procesing across time.

The top-down coupling profile we observed is in accordance with previous results, but we also found more prominent involvement of low frequencies in bottom-up information processing than previously reported. Previous studies found that feedforward (or bottom-up) influences were mediated by gamma frequency range rhythms [Michalareas et al., 2016, Bastos et al., 2015, Fontolan et al., 2014, Bosman et al., 2012]. However, across each of these studies, there were examples of a broader frequency range being employed for bottom-up information transmission. [Fontolan et al., 2014] reported higher bottom-up coupling of delta (1-3Hz) frequency phase in left A1 to high gamma (80-100Hz) amplitude in left association auditory cortex using human depth recordings. [Bastos et al., 2015] found that theta-channels were being used in addition to gamma-frequency channels in feedforward information transmission among primate visual areas. Finally, [Michalareas et al., 2016] did not observe stronger alpha-beta frequency range feedback (top-down) modulation in 8 out of 21 primate visual cortex area pairs (with 2 pairs showing the opposite pattern), and no stronger gamma frequency range feedforward (bottom-up) modulation in 5 out of 21 pairs. We observed dominant bottom-up coupling between low frequency (*θ,α,β*) phase (or amplitude) and low-*γ* amplitude among phase-amplitude couplings (and amplitude-amplitude couplings, respectively). Differences in exact frequency ranges reported may be due to regional (visual versus auditory cortex) variation [Tort et al., 2009, Canolty et al., 2006] and species-related variation. Overall, these findings suggest there is no clear association of particular frequency bands with direction, pointing to a picture of top-down and bottom-up processing that is more complex than the previously hypothesized ‘*β*-down vs. *γ*-up’ [Bastos et al., 2012, Fontolan et al., 2014].

Rather, our findings point to the existence of conserved cross-frequency signatures of bottom-up and top-down information processing in auditory cortex. These patterns revealed a prominent role for low frequencies in mediating both bottom-up and top-down processing, by coupling to different ranges of the target amplitude frequency spectrum. By coupling to *γ* rhythms, low frequency oscillations could shape neuronal excitability directly. High-frequency (e.g. *γ* range) oscillations have been shown to correlate with spiking events. It is also possible for low-frequency oscillations to shape neuronal excitability more indirectly, by coupling to low-frequency rhythms. Through a hierarchically organized oscillation structure, these low-frequency rhythms, in turn, can couple to high-frequency rhythms. Both mechanisms may be at play in regulating information processing in the two directions, but our findings suggest top-down coupling may exert a greater influence over *γ* rhythms.

We observed the same pattern of results in synthetic stimuli. The overall lower coupling strength we observed in synthetic stimuli could be a consequence of the coding differences that were previously observed between synthetic and intact stimuli (Anonymous, 2014), or they could be driving these coding differences. We observed more widespread differences in coupling strength across the frequency spectrum between the two stimulus types among amplitude-amplitude than among phase-amplitude couplings. The *γ* frequency range differences in coupling strength among amplitude-amplitude couplings in particular appear to be more sensitive to acoustic features. Therefore, *γ* range amplitude-amplitude coupling could underlie encoding of stimulus features, whereas phase-amplitude couplings may reflect a more general hallmark of inter-areal communication.

The observation of overall decreased coupling strength in spectrally flat stimuli (EPS) suggests enhanced encoding and processing of information-rich, environmentally relevant stimuli such as the natural vocalization employed here. The fact that envelope-flattened (and spectrally rich) stimuli elicit a coupling profile that resembles natural vocalization stimuli (VOC) more closely suggests some aspects of auditory processing are relatively unhindered by envelope manipulations. This is also consistent with our previous results that showed information about spectrally rich stimuli (SPS) was better maintained than information about spectrally flat stimuli (EPS), at least across the first two sectors (S1,S2) of the auditory hierarchy. In the highest-order auditory sector (S4), decoding performance showed differences across a broader frequency range among VOC and EPS than among VOC and SPS [Anonymous, 2014].

Bottom-up PAC can be relevant to stimulus encoding. For example, an oscillation-based theory of speech encoding [Hyafil et al., 2015a, Giraud and Poeppel, 2012] predicts that cortical theta oscillations track the syllabic rhythm of speech and, in turn, reset the spiking of neurons at a gamma frequency level through *θ*-*γ* PAC. This ensures that neural excitability is optimally aligned with incoming speech. Our finding of dominant low frequency phase to *γ* amplitude coupling bottom-up fits within a framework that requires theta-gamma coupling for accurate speech encoding [Hyafil et al., 2015a]. Cross-regional *θ*-*γ* PAC as we observed here across A1 and higher order sectors could be responsible for coordinating information processing along the auditory cortical hierarchy. Our observation that these patterns are consistent across stimulus classes implies that this may be a generalized mechanism of auditory information processing which is co-opted during speech processing. However, our finding of enhanced low-to-high frequency phase-amplitude coupling in the top-down direction shows these couplings also mediate top-down influences on information processing.

Coupling in the top-down direction has been of particular interest in theories of the effect of attention on auditory information processing. Attention is thought to to sample stimuli rhythmically, fluctuating at a 4-10 Hz rhythm [Landau and Fries, 2012], and was shown to enhance the processing of degraded speech in the anterior and posterior STS [Wild et al., 2012]. Frontal top-down signals in the delta and theta frequency range have been shown to modulate low-frequency oscillations in human auditory cortex, increasing their coupling to continuous speech [Park et al., 2015]. The same delta and theta frequency bands have been shown to be more strongly coupled to gamma-range burst rate within monkey A1 during periods of task entrainment [Lakatos et al., 2016]. In visual cortex, delta frequency oscillations have been shown to entrain with the rhythm of stimuli in the attended stream [Schroeder et al., 2010, Lakatos et al., 2008], and top-down beta-band modulation was found to be enhanced with attention [Bastos et al., 2015]. Increased power in low-frequency range oscillations may also, in turn, enhance bottom-up rhythms through top-down (cross-frequency) coupling [Bressler and Richter, 2015, Lee et al., 2013]; indeed, top-down beta-band activity was found to be maximally correlated with bottom-up gamma-band activity when top-down preceded bottom-up activity [Richter et al., 2017]. In addition, top-down couplings may also reflect the modulatory activity of predictive processes on auditory/speech perception & processing [Chao et al., 2018, Arnal and Giraud, 2012, Gagnepain et al., 2012, Davis et al., 2011, Wild et al., 2010, Poeppel et al., 2008]. Our findings of dominant low-to-low and low-to-high frequency coupling are consistent with these accounts, and could well serve to enhance the animals engagement in the task [Lakatos et al., 2016, Park et al., 2015]. In addition, the top-down coupling signal we observed may also be carrying predictive information, being consistent with frequency ranges which have been shown to carry prediction-update signals [Chao et al., 2018]. We cannot dissociate between attentive or predictive signals here, however.

We found PAC and AAC to offer similar accounts of information processing. This could be due to a high degree of shared informational content between the two measures; it remains unclear how these measures differ, and how AAC could function mechanistically in the brain. To the degree that PAC and AAC offer complementary, non-overlapping information, it is possible that top-down influences are exerted more broadly through various CFC types including both PAC and AAC, and others such as phase synchronization [Fries, 2005].

Comparisons of bottom-up and top-down coupling strength are relevant to models of neural information processing such as predictive coding. We found that low frequencies were prominently involved in both bottom-up and top-down information processing through cross-frequency coupling. Thus the prediction of *highfrequencies*(*γ*)*up, lowfrequencies*(*β*)*down* does not generalize to cross-frequency interactions. Rather, we found preserved signatures for bottom-up and top-down information processing among cross-frequency interactions, in which low frequencies predominantly targeted different *γ* amplitude ranges in the two directions. Within a cross-frequency coupling framework, prediction errors and predictions may thus be propagated up and down the cortical hierarchy, respectively, through PAC: low frequency (*θ,α,β*) phase to *highγ* couplings could be used mainly to propagate predictions top-down, adjusting lower-order receptivity according to expectations. Meanwhile, prediction errors could be relayed to higher order areas through cross-regional low frequency-to-*lowγ* interactions. Also, information could be relayed using the same frequency ranges by alternating coupling across time, as we have observed among low frequency PAC. Future studies may establish whether (and to what extent) cross-frequency interactions carry prediction and prediction error signals.

In summary, we revealed distinct cross-frequency signatures of top-down and bottom-up information processing. These signatures are largely preserved across coupling types and stimulus types, suggesting they are a general hallmark of information processing in auditory cortex. Our findings suggest that information need not be propagated along separate frequency channels up and down the auditory cortical hierarchy. Rather, low frequencies prominently featured in both bottom-up and top-down information processing across multiple auditory sectors in macaques. Our finding of prominent low-to-high frequency coupling top-down extends the current view of theta-gamma interactions in auditory cortex, which primarily emphasized their involvement in bottom-up processes. Moreover, our findings suggest low frequencies may be used for bi-directional information processing by coupling to distinct frequency ranges in the target region, and by alternating bottom-up and top-down processing across time. This extends our understanding of how information is propagated up and down the cortical hierarchy, with significant implications for theories of predictive coding.

## 6. Supporting information

**Fig S1.**
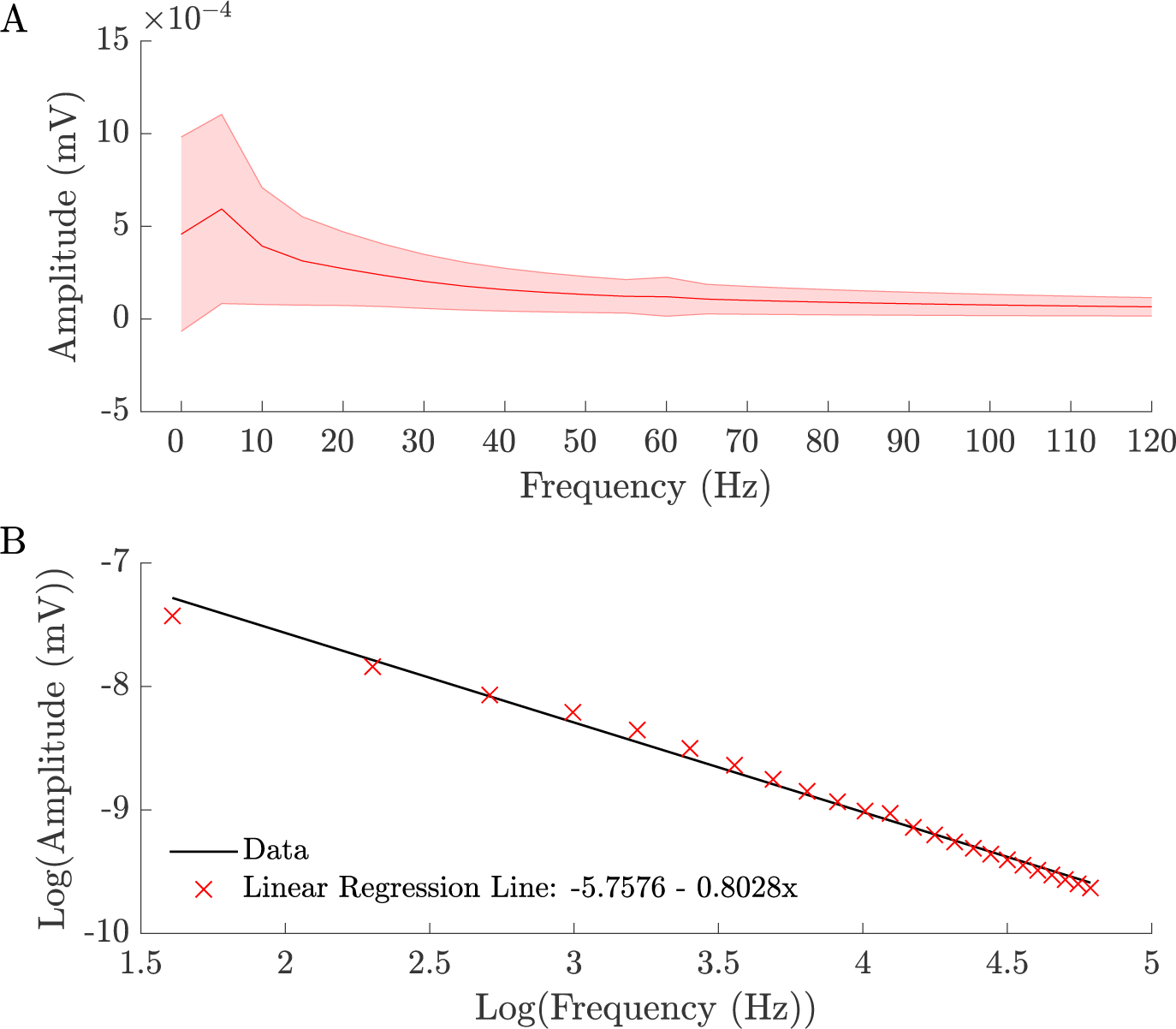
(A) Power vs. frequency of neural signal and (B) regression line fit. The amplitude of the neural recording signal peaks in the *α* (7.5-12.5 Hz) frequency range and shows the typical 1/f decay pattern thereafter. The signal depicted here is from stimulus onset (time=0) for presentation of the intact stimulus (VOC), averaged across all variables. Standard deviation across all variables is depicted by the red shaded region. The regression line fit to the neural signal confirms a 1/f decay pattern (B). The *δ* frequency band (0-2.5 Hz) was excluded from analyses (see Methods and Materials).

**Fig S2.**
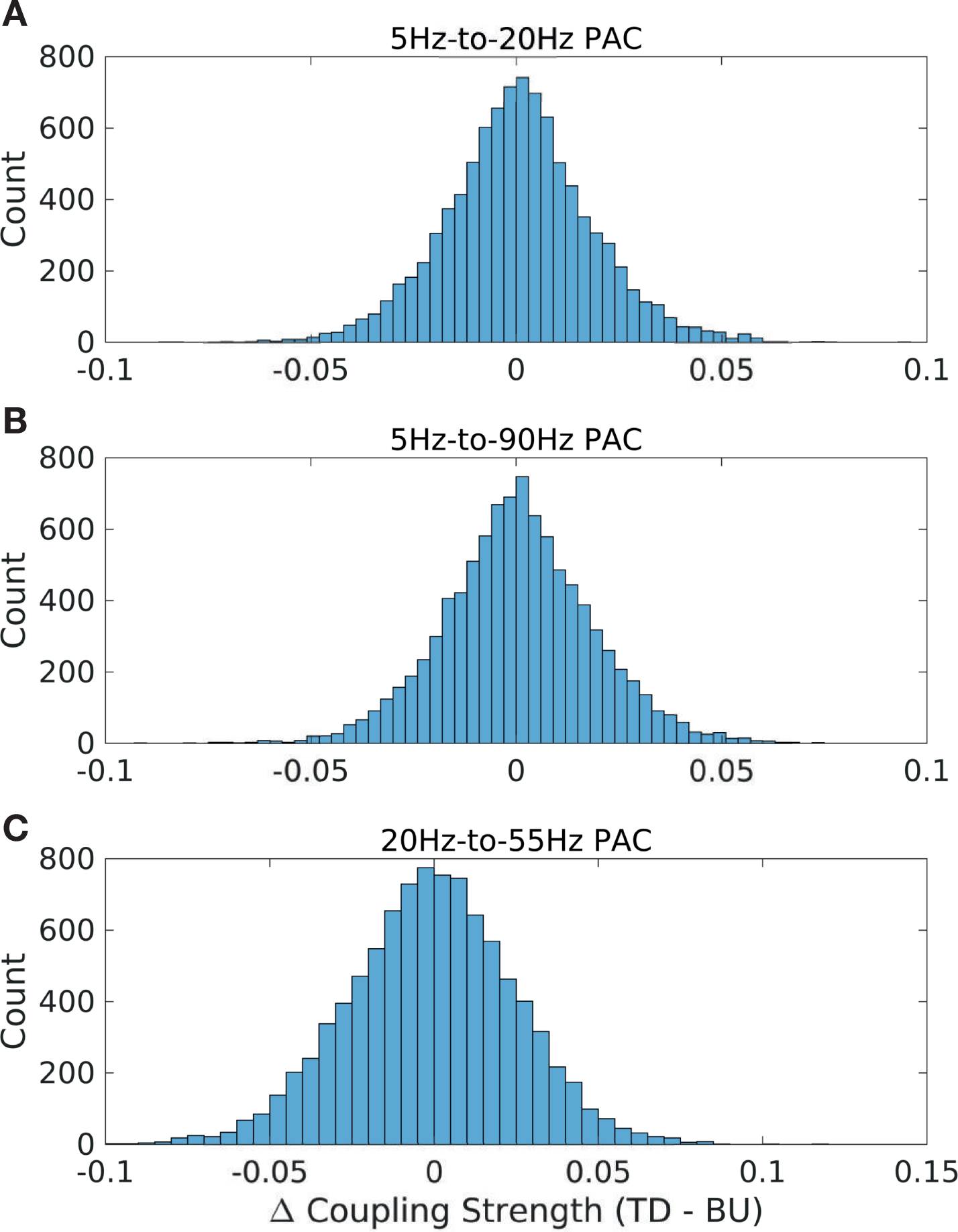
Distribution of differences in PAC coupling strength for select CFC pairs in natural stimuli (VOC) **(A)** *θ* (5Hz) to *β* (20Hz) PAC **(B)** *θ* (5Hz) to *highγ* (90Hz) PAC **(C)** *β* (20Hz) to *lowγ* (55Hz) PAC

**Fig S3.**
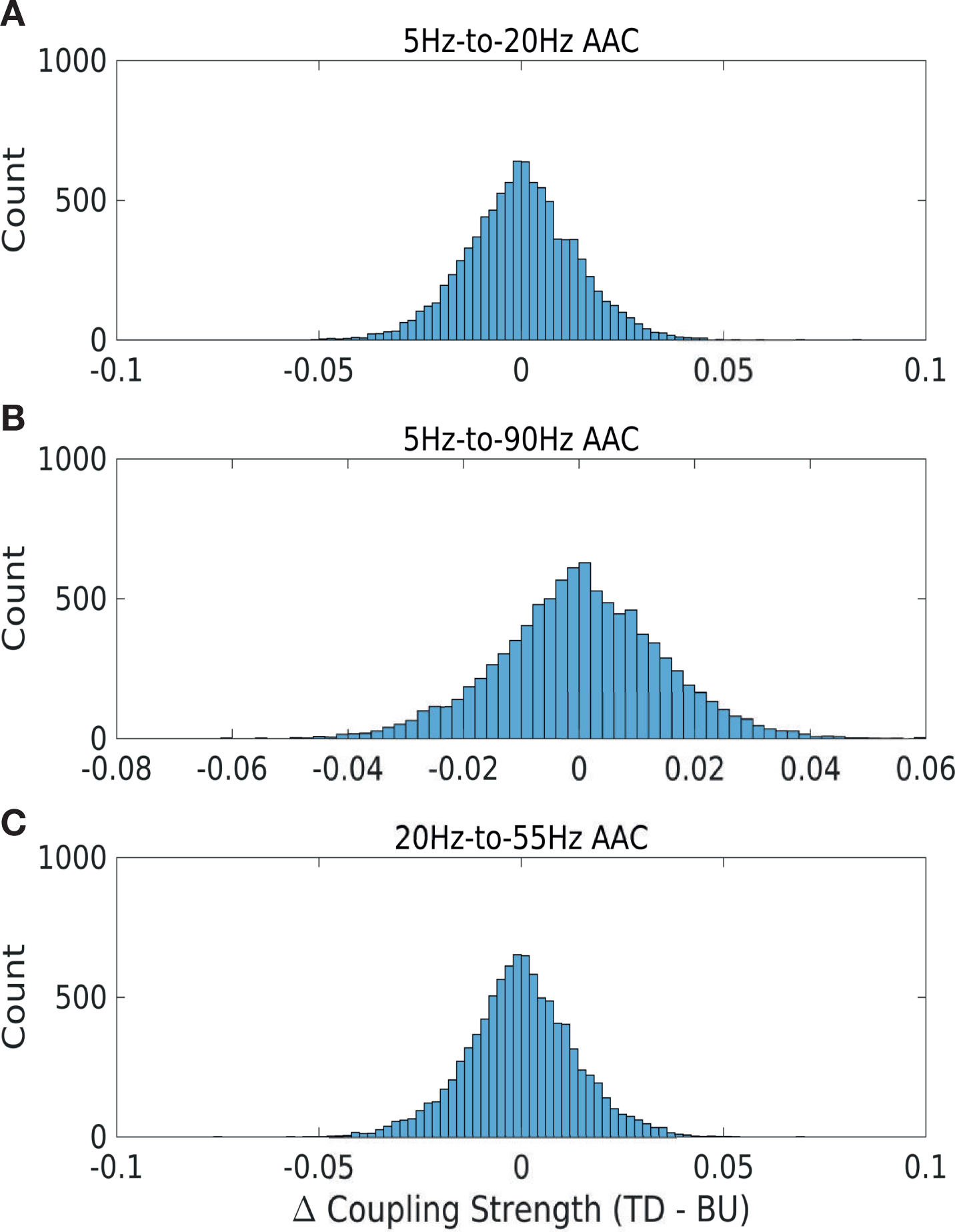
Distribution of differences in AAC coupling strength for select CFC pairs in natural stimuli (VOC) **(A)** *θ* (5Hz) to *β* (20Hz) PAC **(B)** *θ* (5Hz) to *highγ* (90Hz) PAC **(C)** *β* (20Hz) to *lowγ* (55Hz) PAC

**Fig S4.**
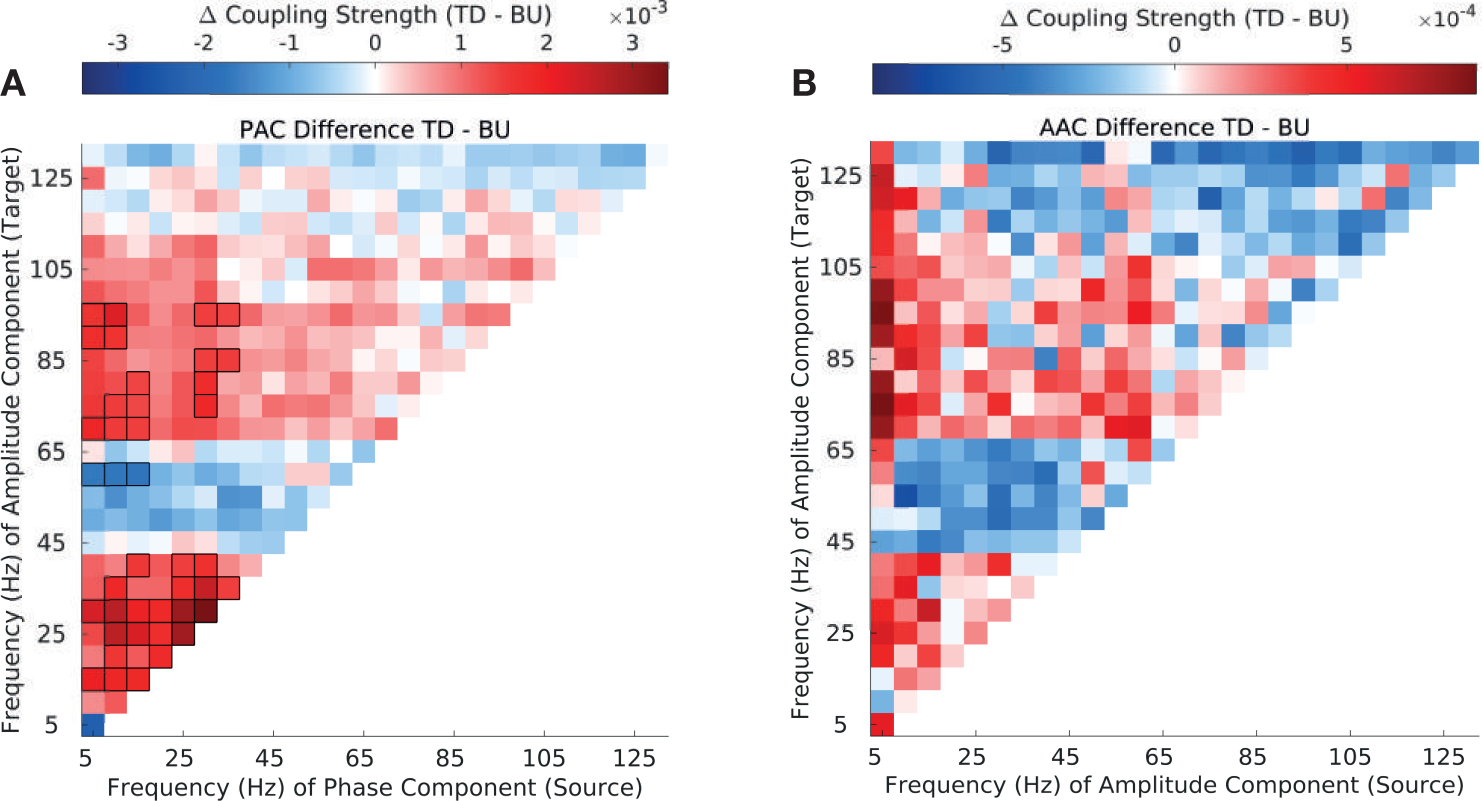
Differences in top-down and bottom-up PAC and AAC coupling strength in natural stimuli (VOC) with significance via cluster-permutation test. **(A)** Difference in top-down and bottom-up in PAC strength. Significant differences are enclosed by black rectangles (cluster-permutation test, p-value=0.01). **(B)** Difference between top-down and bottom-up in AAC strength. Significant differences are enclosed by black rectangles (cluster-permutation test, p-value=0.01). Results are depicted averaged across all channels, cross-regional pairs, and animals.

**Fig S5.**
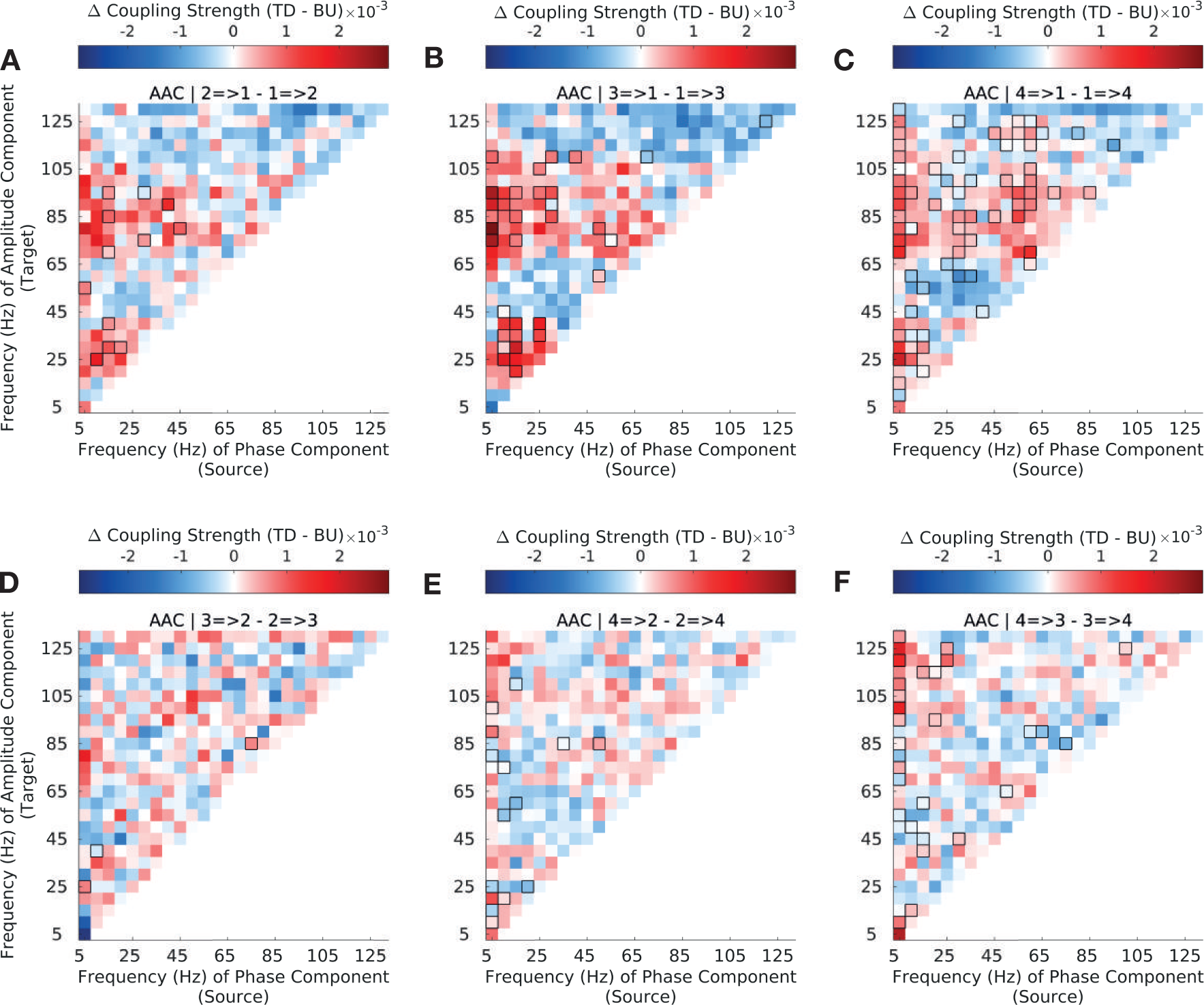
Top-down versus bottom-up amplitude-amplitude coupling strength (AAC) across the auditory hierarchy in natural stimuli (VOC) Difference in top-down and bottom-up *amplitude-amplitude* coupling (AAC) strength for CFC between sectors 1 (A1/ML) and 2 (R/AL) **(A)**, sectors 1 (A1/ML) and 3 (RTL) **(B)**, sectors 1 (A1/ML) and 4 (RTp) **(C)**, sectors 2 and 3 **(D)**, sectors 2 and 4 **(E)** and sectors 3 and 4 **(F)** (see Methods and Materials 3.5 for sector definitions). Interactions between sector 1 (A1/ML) and higher order sectors show strong asymmetries in bottom-up and top-down coupling strength across the frequency spectrum; interactions among higher-order sectors (S2-S4) show less widespread asymmetries. Significant differences are enclosed by black rectangles (FDR corrected, q-value=0.01). Results are depicted averaged across all channels and animals.

**Fig S6.**
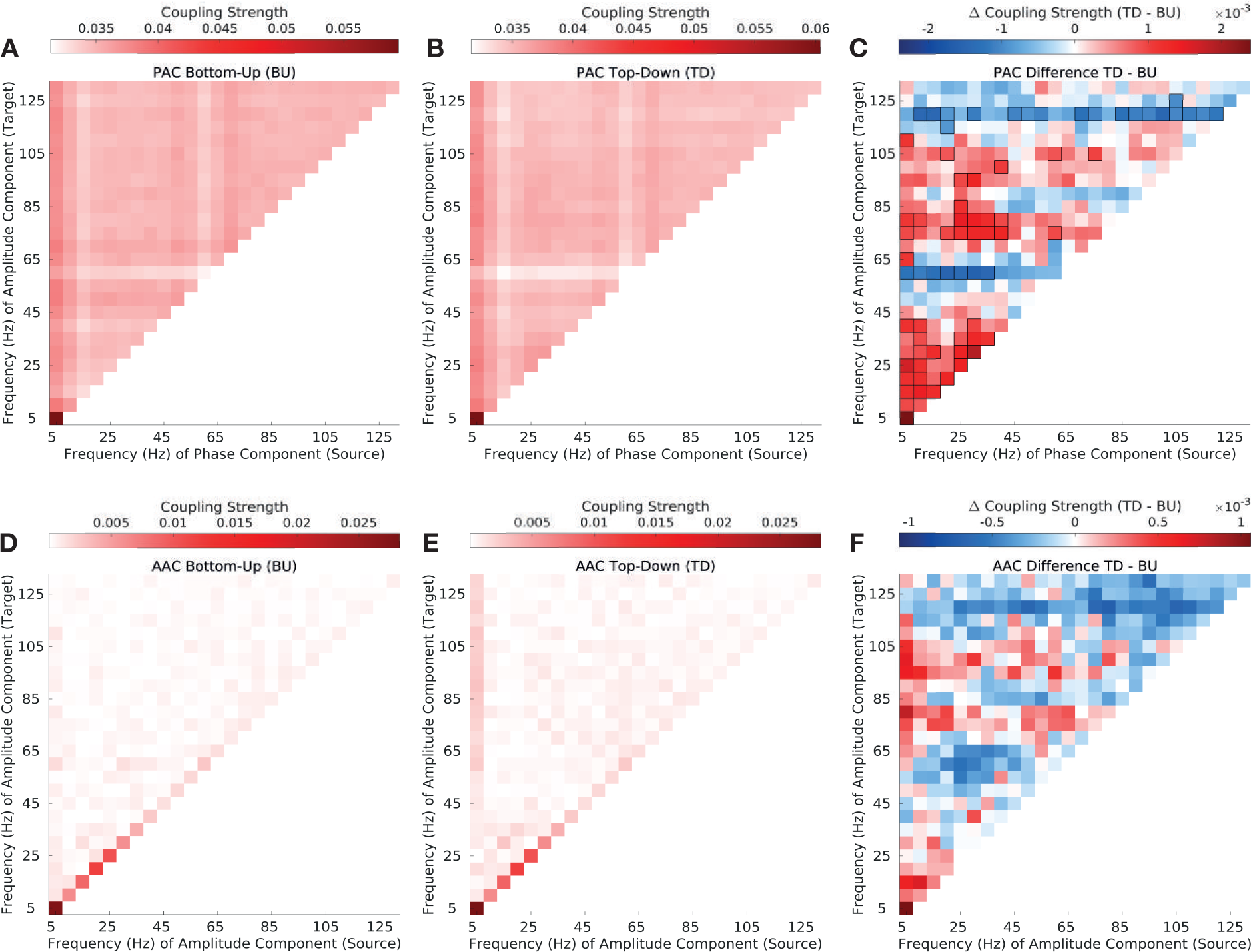
Top-down vs. bottom-up phase-amplitude and amplitude-amplitude coupling in synthetic envelope-preserved vocalizations (EPS). **(A-B)** Phase-amplitude coupling (PAC) strength in the top-down (A) and bottom-up direction (B). Depicted are canonical correlation-derived coupling coefficients (see Methods and Materials). **(C)** Difference in top-down and bottom-up in PAC strength. Significant differences are enclosed by black rectangles (FDR corrected, q-value=0.01). **(D-E)** Amplitude-amplitude coupling (AAC) strength in the top-down (D) and bottom-up direction (E). Depicted are canonical correlation-derived coupling coefficients. **(F)** Difference between top-down and bottom-up in AAC strength. Significant differences are enclosed by black rectangles (FDR corrected, q-value=0.01). Results are depicted averaged across all channels, cross-regional pairs, and animals.

**Fig S7.**
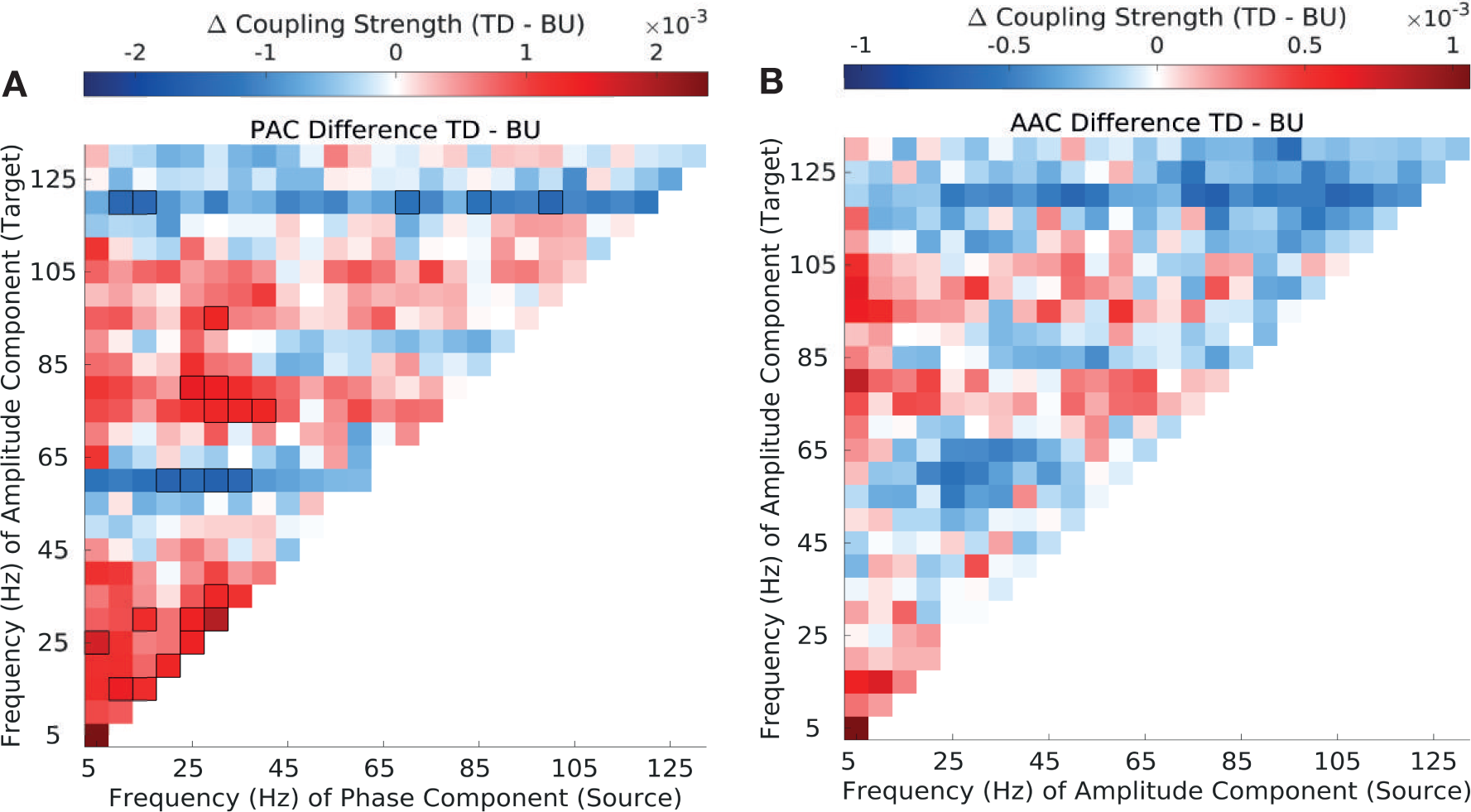
Differences in top-down and bottom-up PAC and AAC coupling strength in synthetic envelope-preserved stimuli (EPS) with significance via cluster-permutation test. **(A)** Difference in top-down and bottom-up in PAC strength. Significant differences are enclosed by black rectangles (cluster-permutation test, p-value=0.01). **(B)** Difference between top-down and bottom-up in AAC strength. Significant differences are enclosed by black rectangles (cluster-permutation test, p-value=0.01). Results are depicted averaged across all channels, cross-regional pairs, and animals.

**Fig S8.**
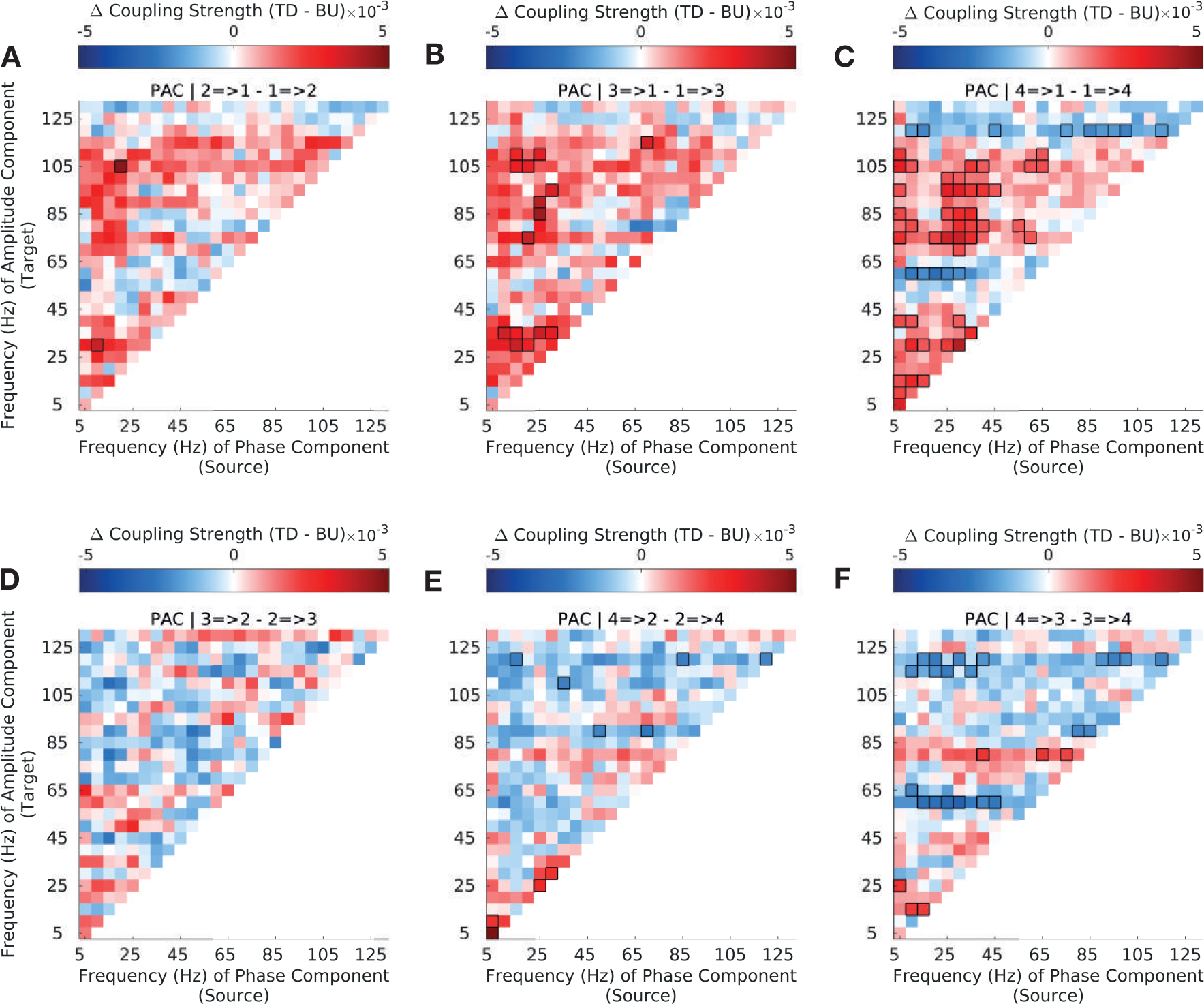
Top-down versus bottom-up phase-amplitude coupling strength (PAC) across the auditory hierarchy in synthetic envelop-preserved stimuli (EPS) Difference in top-down and bottom-up *phase-amplitude* coupling (PAC) strength for CFC between sectors 1 (A1/ML) and 2 (R/AL) **(A)**, sectors 1 (A1/ML) and 3 (RTL) **(B)**, sectors 1 (A1/ML) and 4 (RTp) **(C)**, sectors 2 and 3 **(D)**, sectors 2 and 4 **(E)** and sectors 3 and 4 **(F)** (see Methods and Materials 3.5 for sector definitions). Interactions between sector 1 (A1/ML) and higher order sectors show strong asymmetries in bottom-up and top-down coupling strength across the frequency spectrum; interactions among higher-order sectors (S2-S4) show less widespread asymmetries. Significant differences are enclosed by black rectangles (FDR corrected, q-value=0.01). Results are depicted averaged across all channels and animals.

**Fig S9.**
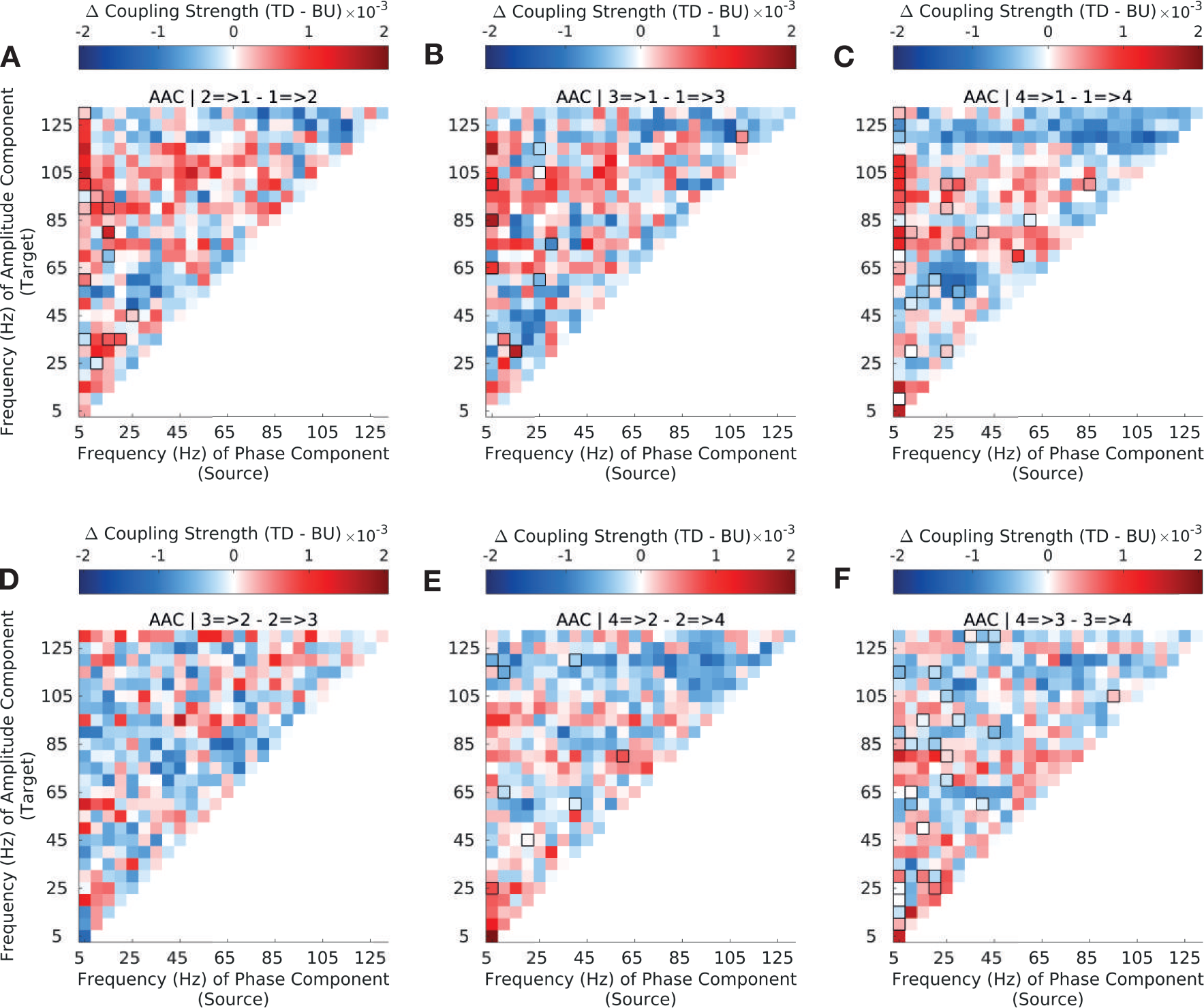
Top-down versus bottom-up amplitude-amplitude coupling strength (AAC) across the auditory hierarchy in synthetic envelope-preserved stimuli (EPS) Difference in top-down and bottom-up *amplitude-amplitude* coupling (AAC) strength for CFC between sectors 1 (A1/ML) and 2 (R/AL) **(A)**, sectors 1 (A1/ML) and 3 (RTL) **(B)**, sectors 1 (A1/ML) and 4 (RTp) **(C)**, sectors 2 and 3 **(D)**, sectors 2 and 4 **(E)** and sectors 3 and 4 **(F)** (see Methods and Materials 3.5 for sector definitions). Interactions between sector 1 (A1/ML) and higher order sectors show strong asymmetries in bottom-up and top-down coupling strength across the frequency spectrum; interactions among higher-order sectors (S2-S4) show less widespread asymmetries. Significant differences are enclosed by black rectangles (FDR corrected, q-value=0.01). Results are depicted averaged across all channels and animals.

**Fig S10.**
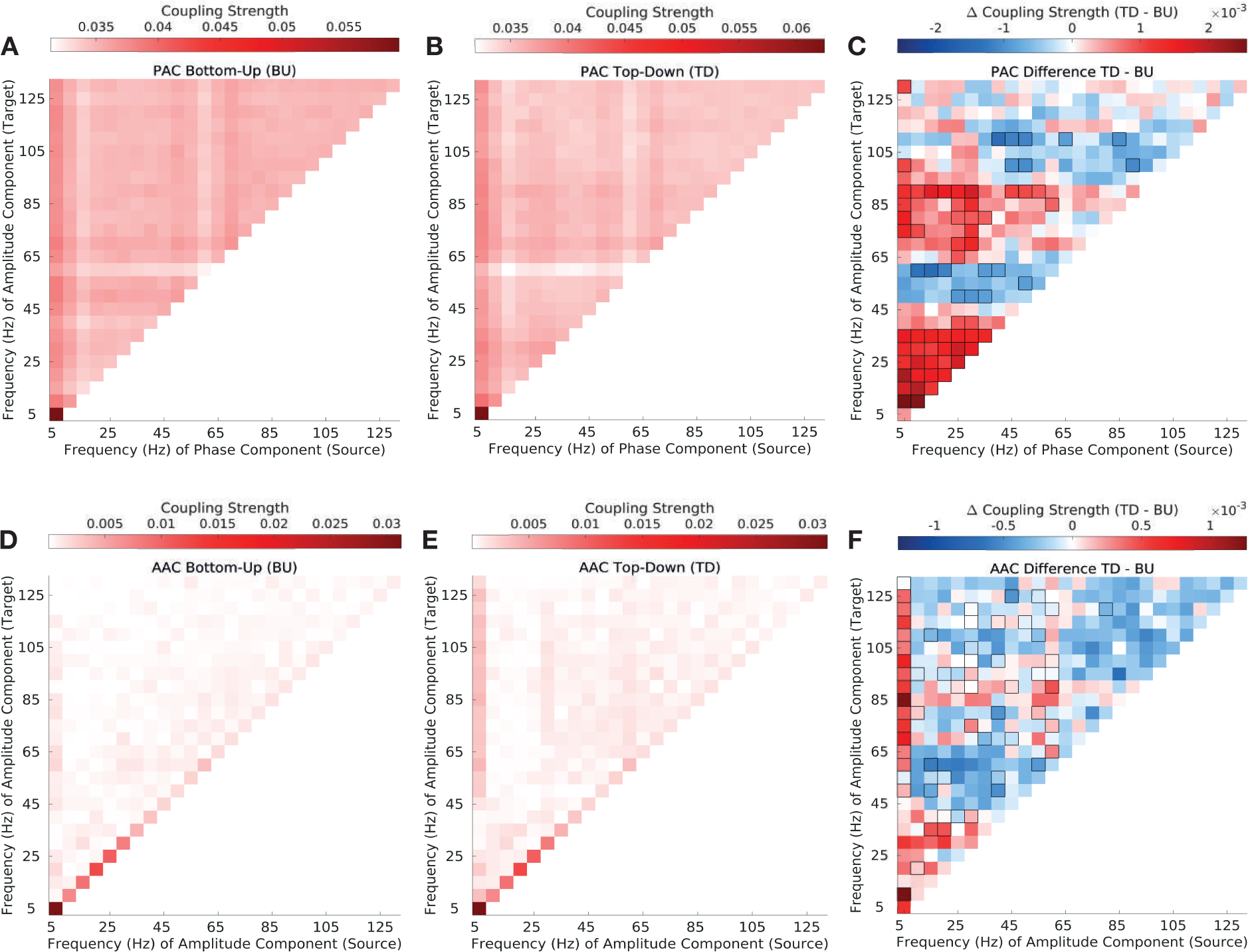
Top-down vs. bottom-up phase-amplitude and amplitude-amplitude coupling in synthetic spectrum-preserved vocalizations (SPS). **(A-B)** Phase-amplitude coupling (PAC) strength in the top-down (A) and bottom-up direction (B). Depicted are canonical correlation-derived coupling coefficients (see Methods and Materials). **(C)** Difference in top-down and bottom-up in PAC strength. Significant differences are enclosed by black rectangles (FDR corrected, q-value=0.01). **(D-E)** Amplitude-amplitude coupling (AAC) strength in the top-down (D) and bottom-up direction (E). Depicted are canonical correlation-derived coupling coefficients. **(F)** Difference between top-down and bottom-up in AAC strength. Significant differences are enclosed by black rectangles (FDR corrected, q-value=0.01). Results are depicted averaged across all channels, cross-regional pairs, and animals.

**Fig S11.**
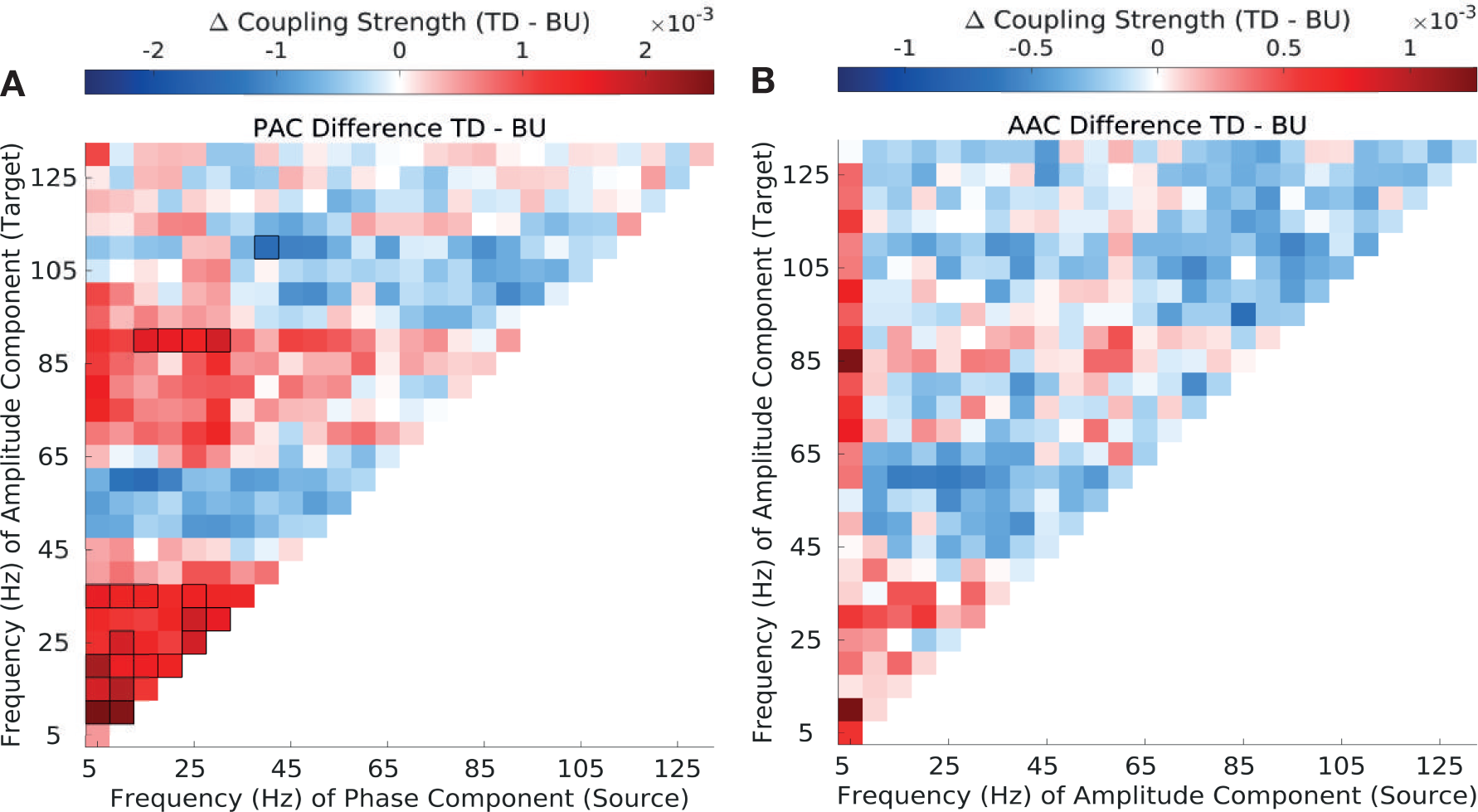
Differences in top-down and bottom-up PAC and AAC coupling strength in synthetic spectrum-preserved stimuli (SPS) with significance via cluster-permutation test. **(A)** Difference in top-down and bottom-up in PAC strength. Significant differences are enclosed by black rectangles (cluster-permutation test, p-value=0.01). **(B)** Difference between top-down and bottom-up in AAC strength. Significant differences are enclosed by black rectangles (cluster-permutation test, p-value=0.01). Results are depicted averaged across all channels, cross-regional pairs, and animals.

**Fig S12.**
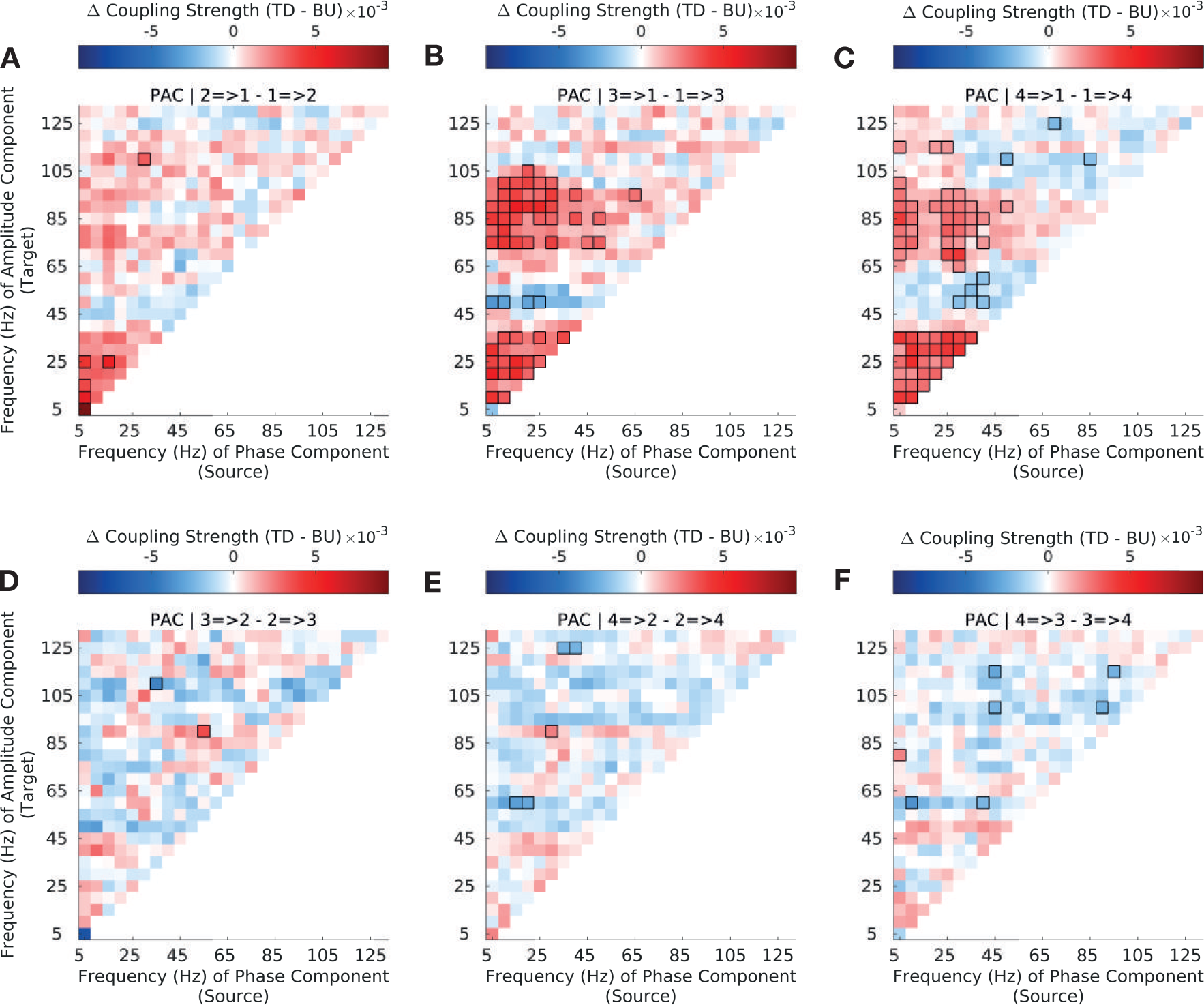
Top-down versus bottom-up phase-amplitude coupling strength (PAC) across the auditory hierarchy in synthetic spectrum-preserved stimuli (SPS) Difference in top-down and bottom-up *phase-amplitude* coupling (PAC) strength for CFC between sectors 1 (A1/ML) and 2 (R/AL) **(A)**, sectors 1 (A1/ML) and 3 (RTL) **(B)**, sectors 1 (A1/ML) and 4 (RTp) **(C)**, sectors 2 and 3 **(D)**, sectors 2 and 4 **(E)** and sectors 3 and 4 **(F)** (see Methods and Materials 3.5 for sector definitions). Interactions between sector 1 (A1/ML) and higher order sectors show strong asymmetries in bottom-up and top-down coupling strength across the frequency spectrum; interactions among higher-order sectors (S2-S4) show less widespread asymmetries. Significant differences are enclosed by black rectangles (FDR corrected, q-value=0.01). Results are depicted averaged across all channels and animals.

**Fig S13.**
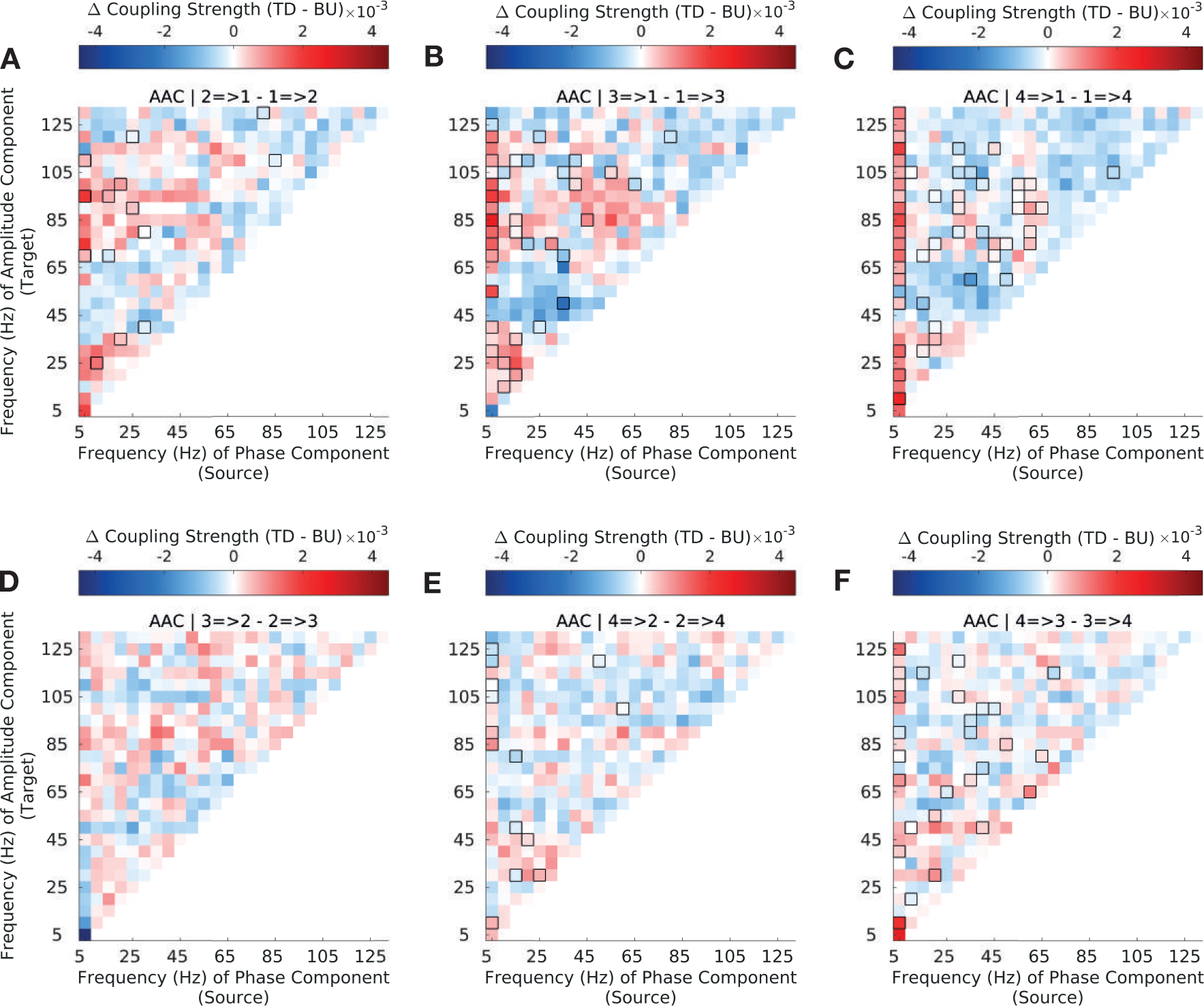
Top-down versus bottom-up amplitude-amplitude coupling strength (AAC) across the auditory hierarchy in synthetic spectrum-preserved stimuli (SPS) Difference in top-down and bottom-up *amplitude-amplitude* coupling (AAC) strength for CFC between sectors 1 (A1/ML) and 2 (R/AL) **(A)**, sectors 1 (A1/ML) and 3 (RTL) **(B)**, sectors 1 (A1/ML) and 4 (RTp) **(C)**, sectors 2 and 3 **(D)**, sectors 2 and 4 **(E)** and sectors 3 and 4 **(F)** (see Methods and Materials 3.5 for sector definitions). Interactions between sector 1 (A1/ML) and higher order sectors show strong asymmetries in bottom-up and top-down coupling strength across the frequency spectrum; interactions among higher-order sectors (S2-S4) show less widespread asymmetries. Significant differences are enclosed by black rectangles (FDR corrected, q-value=0.01). Results are depicted averaged across all channels and animals.

**Fig S14.**
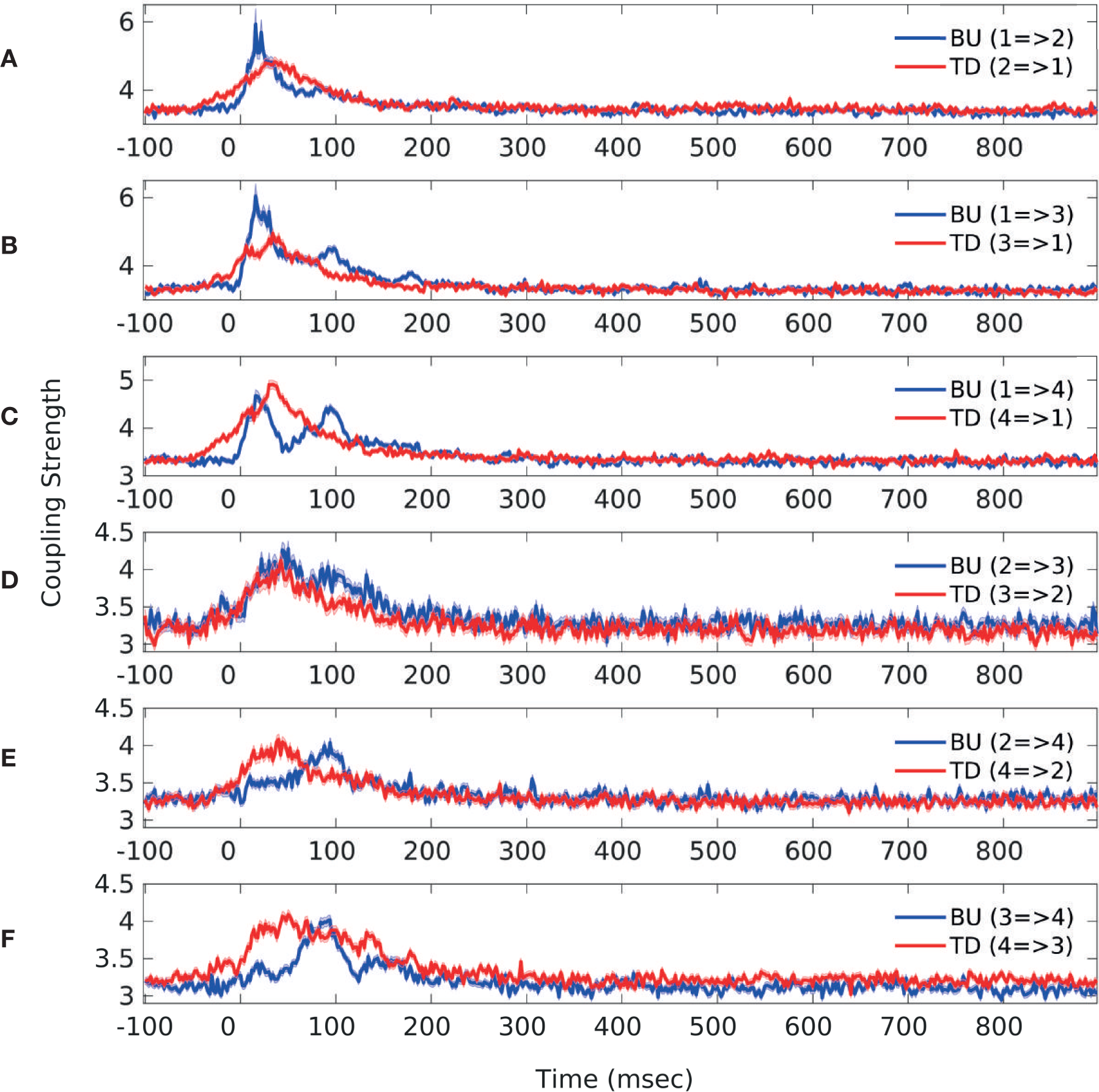
Top-down vs. bottom-up low-frequency PAC over time across the auditory hierarchy. Low frequency phase (*θ*, 2.5-7.5Hz) to low frequency amplitude (*β*, 15-25Hz) coupling over the duration of stimulus presentation, separately in the bottom-up direction (blue) and the top-down direction (red), depicted for all cross-regional pairs. **A** S2=>S1 (red), S1=>S2 (blue) **B** S3=>S1 (red), S1=>S3 (blue) **C** S4=>S1 (red), S1=>S4 (blue) **D** S3=>S2 (red), S2=>S3 (blue) **E** S4=>S2 (red), S2=>S4 (blue) **F** S4=>S3 (red), S3=>S4 (blue)

**Fig S15.**
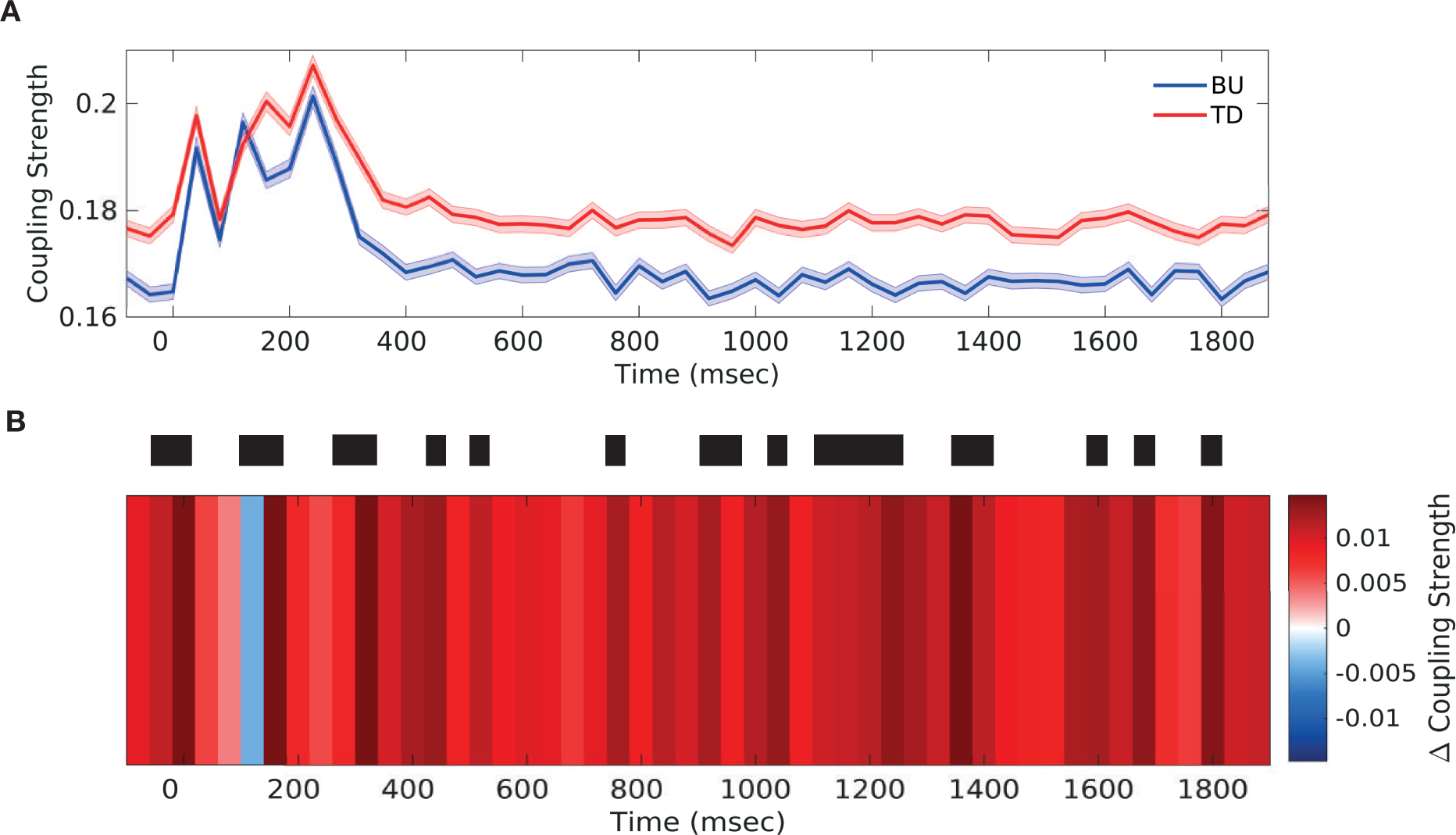
Top-down vs. bottom-up low-frequency PAC over time computed using short-time fast Fourier Transform (FFT) **(A)** Low frequency phase (*θ*/*α*, 2.5-12.5Hz) to low frequency amplitude (*β*, 12.5-37.5Hz) coupling over the duration of stimulus presentation, separately in the bottom-up direction (blue) and the top-down direction (red) **(B)** Differences in top-down and bottom-up low frequency PAC (*θ*/*α*, (2.5-12.5Hz) phase to *β* (12.5-37.5Hz) amplitude coupling) over the duration of stimulus presentation. Significant differences are marked by thick black lines (FDR corrected, q-value=0.01). Results are depicted averaged across all channels, cross-regional pairs, and animals. Top-down and bottom-up coupling were found to alternate in the period around stimulus onset (−50ms to 200ms).

## References

Arnal, L. H. and Giraud, A.-L. (2012). Cortical oscillations and sensory predictions. Trends in Cognitive Sciences, 16(7):390–398.

Aru, J., Aru, J., Priesemann, V., Wibral, M., Lana, L., Pipa, G., Singer, W., and Vicente, R. (2015). Untangling cross-frequency coupling in neuroscience. Current Opinion in Neurobiology, 31:51–61.

Bastos, A. M., Usrey, W. M., Adams, R. A., Mangun, G. R., Fries, P., and Friston, K. J. (2012). Canonical Microcircuits for Predictive Coding. Neuron, 76(4):695–711.

Bastos, A. M., Vezoli, J., Bosman, C. A., Schoffelen, J.-M., Oostenveld, R., Dowdall, J. R., De Weerd, P., Kennedy, H., and Fries, P. (2015). Visual Areas Exert Feedforward and Feedback Influences through Distinct Frequency Channels. Neuron, 85(2):390–401.

Bosman, C. A., Schoffelen, J.-M., Brunet, N., Oostenveld, R., Bastos, A. M., Womelsdorf, T., Rubehn, B., Stieglitz, T., De Weerd, P., and Fries, P. (2012). Attentional Stimulus Selection through Selective Synchronization between Monkey Visual Areas. Neuron, 75(5):875–888.

Bressler, S. L. and Richter, C. G. (2015). ScienceDirect Interareal oscillatory synchronization in top-down neocortical processing. Current Opinion in Neurobiology, 31:62–66.

Buzsáaki, G., Anastassiou, C. A., and Koch, C. (2012). The origin of extracellular fields and currents — EEG, ECoG, LFP and spikes. Nature Reviews Neuroscience, 13(6):1–14.

Canolty, R. T., Edwards, E., Dalal, S. S., Soltani, M., Nagarajan, S. S., Kirsch, H. E., Berger, M. S., Barbaro, N. M., and Knight, R. T. (2006). High Gamma Power Is Phase-Locked to Theta Oscillations in Human Neocortex. Science, 313(5793):1626–1628.

Canolty, R. T. and Knight, R. T. (2010). The functional role of cross-frequency coupling. Trends in Cognitive Sciences, 14(11):506–515.

Chao, Z. C., Takaura, K., Wang, L., Fujii, N., and Dehaene, S. (2018). Large-Scale Cortical Networks for Hierarchical Prediction and Prediction Error in the Primate Brain. Neuron, pages 1–30.

Cohen, M. X. (2008). Assessing transient cross-frequency coupling in EEG data. Journal of Neuroscience Methods, 168(2):494–499.

Colgin, L. L. (2015). Theta–gamma coupling in the entorhinal–hippocampal system. Current Opinion in Neurobiology, 31:45–50.

Davis, M. H., Ford, M. A., Kherif, F., and Johnsrude, I. S. (2011). Does Semantic Context Benefit Speech Understanding through. Journal of Cognitive Neuroscience, pages 1–21.

Doesburg, S. M., Green, J. J., McDonald, J. J., and Ward, L. M. (2012). Theta modulation of inter-regional gamma synchronization during auditory attention control. Brain Research, 1431(C):77–85.

Durstewitz, D. (2017). Advanced Data Analysis in Neuroscience. Springer International Publishing.

Fontolan, L., Morillon, B., Liegeois-Chauvel, C., and Giraud, A.-L. (2014). The contribution of frequency-specific activity to hierarchical information processing in the human auditory cortex. Nature Communications, 5(1):1–10.

Fries, P. (2005). A mechanism for cognitive dynamics: neuronal communication through neuronal coherence. Trends in Cognitive Sciences, 9(10):474–480.

Friston, K. (2008). Hierarchical Models in the Brain. PLOS Computational Biology, 4(11):e1000211–24.

Gagnepain, P., Henson, R. N., and Davis, M. H. (2012). Temporal Predictive Codes for Spoken Words in Auditory Cortex. Current Biology, 22(7):615–621.

Giraud, A.-L. and Poeppel, D. (2012). Cortical oscillations and speech processing: emerging computational principles and operations. Nature Neuroscience, 15(4):511–517.

Hyafil, A., Fontolan, L., Kabdebon, C., Gutkin, B., and Giraud, A.-L. (2015a). Speech encoding by coupled cortical theta and gamma oscillations. eLife, 4:3958.

Hyafil, A., Giraud, A.-L., Fontolan, L., and Gutkin, B. (2015b). Neural Cross-Frequency Coupling: Connecting Architectures, Mechanisms, and Functions. Trends in Neurosciences, 38(11):725–740.

Jensen, O. and Colgin, L. L. (2007). Cross-frequency coupling between neuronal oscillations. Trends in Cognitive Sciences, 11(7):267–269.

Jirsa, V. and Mueller, V. (2013). Cross-frequency coupling in real and virtual brain networks. Frontiers in Computational Neuroscience, pages 1–25.

Kellis, S., Miller, K., Thomson, K., Brown, R., House, P., and Greger, B. (2010). Decoding spoken words using local field potentials recorded from the cortical surface. Journal of Neural Engineering, 7(5):056007–20.

Kendrick, K. M., Zhan, Y., Fischer, H., Nicol, A. U., Zhang, X., and Feng, J. (2011). Learning alters theta amplitude, theta-gamma coupling and neuronal synchronization in inferotemporal cortex. BMC Neuroscience, 12(1):55.

Kikuchi, Y., Horwitz, B., and Mishkin, M. (2010). Hierarchical auditory processing directed rostrally along the monkey’s supratemporal plane. The Journal of neuroscience : the official journal of the Society for Neuroscience, 30(39):13021–13030.

Kveraga, K., Ghuman, A. S., and Bar, M. (2007). Top-down predictions in the cognitive brain. Brain and Cognition, 65(2):145–168.

Lakatos, P., Barczak, A., Neymotin, S. A., McGinnis, T., Ross, D., Javitt, D. C., and O’Connell, M. N. (2016). Global dynamics of selective attention and its lapses in primary auditory cortex. Nature Neuroscience, 19(12):1707–1717.

Lakatos, P., Karmos, G., Mehta, A. D., Ulbert, I., and Schroeder, C. E. (2008). Entrainment of Neuronal Oscillations as a Mechanism of Attentional Selection. Science, 320(5872):110–113.

Landau, A. N. and Fries, P. (2012). Attention Samples Stimuli Rhythmically. Current Biology, 22(11):1000–1004.

Lee, J. H., Whittington, M. A., and Kopell, N. J. (2013). Top-Down Beta Rhythms Support Selective Attention via Interlaminar Interaction: A Model. PLOS Computational Biology, 9(8):e1003164–23.

Maris, E. and Oostenveld, R. (2007). Nonparametric statistical testing of EEG- and MEG-data. Journal of Neuroscience Methods, 164(1):177–190.

Michalareas, G., Vezoli, J., van Pelt, S., Schoffelen, J.-M., Kennedy, H., and Fries, P. (2016). Alpha-Beta and Gamma Rhythms Subserve Feedback and Feedforward Influences among Human Visual Cortical Areas. Neuron, 89(2):384–397.

Park, H., Ince, R. A. A., Schyns, P. G., Thut, G., and Gross, J. (2015). Frontal Top-Down Signals Increase Coupling of Auditory Low-Frequency Oscillations to Continuous Speech in Human Listeners. CURBIO, 25(12):1649–1653.

Poeppel, D., Idsardi, W. J., and van Wassenhove, V. (2008). Speech perception at the interface of neurobiology and linguistics. Philosophical Transactions of the Royal Society B: Biological Sciences, 363(1493):1071–1086.

Ray, S. and Maunsell, J. H. R. (2011). Different Origins of Gamma Rhythm and High-Gamma Activity in Macaque Visual Cortex. PLoS Biology, 9(4):e1000610–15.

Richter, C. G., Thompson, W. H., Bosman, C. A., and Fries, P. (2017). Top-Down Beta Enhances Bottom-Up Gamma. Journal of Neuroscience, 37(28):6698–6711.

Romanski, L. M. and Averbeck, B. B. (2009). The Primate Cortical Auditory System and Neural Representation of Conspecific Vocalizations. Annual Review of Neuroscience, 32(1):315–346.

Salin, P.-A. and Bullier, J. (1995). Corticocortical Connections in the Visual System: Structure and Function. Physiological Review, 75(1):1–48.

Schack, B., Vath, N., Petsche, H., Geissler, H. G., and Moeller, E. (2002). Phase-coupling of theta-gamma EEG rhythms during short-term memory processing. International Journal of Psychophysiology, 44:143–163.

Schroeder, C. E., Wilson, D. A., Radman, T., Scharfman, H., and Lakatos, P. (2010). Dynamics of Active Sensing and perceptual selection. Current Opinion in Neurobiology, 20(2):172–176.

Scott, B. H., Leccese, P. A., Saleem, K. S., Kikuchi, Y., Mullarkey, M. P., Fukushima, M., Mishkin, M., and Saunders, R. C. (2015). Intrinsic Connections of the Core Auditory Cortical Regions and Rostral Supratemporal Plane in the Macaque Monkey. Cerebral cortex (New York, N.Y. : 1991), 7:bhv277–32.

Tort, A. B. L., Komorowski, R. W., Manns, J. R., Kopell, N. J., and Eichenbaum, H. (2009). Theta–gamma coupling increases during the learning of item–context associations. PNAS, 106(49):20942–20947.

Tort, A. B. L., Kramer, M. A., Thorn, C., Gibson, D. J., Kubota, Y., Graybriel, A. M., and Kopell, N. J. (2008). Dynamic cross-frequency couplings of local field potential oscillations in rat striatum and hippocampus during performance of a T-maze task. PNAS, 105(51):20517–20522.

Wild, C. J., Davis, M. H., and Johnsrude, I. S. (2010). Human auditory cortex is sensitive to the perceived clarity of speech. NeuroImage, 60(2):1–13.

Wild, C. J., Yusuf, A., Wilson, D. E., Peelle, J. E., Davis, M. H., and Johnsrude, I. S. (2012). Effortful Listening: The Processing of Degraded Speech Depends Critically on Attention. Journal of Neuroscience, 32(40):14010–14021.

